# Histone H4 lysine 20 monomethylation is not a mark of transcriptional silencers

**DOI:** 10.1101/2025.01.09.632211

**Authors:** Julian A. Segert, Nikola A. Mizgier, Karen Adelman, Martha L. Bulyk

## Abstract

Transcriptional silencers are *cis*-regulatory elements that downregulate the expression of target genes. Although thousands of silencers have been identified experimentally, a predictive chromatin signature of silencers has not been found. H4K20me1 previously was reported to be highly enriched among human silencers, but our reanalysis of the data using an appropriate background revealed that the enrichment is only marginal. We generated H4K20me1 ChIP-seq profiles in *Drosophila* S2 cells, which similarly showed that H4K20me1 does not mark *Drosophila* silencers and instead is associated with active transcription. Silencers remain a poorly annotated, difficult to predict class of *cis*-regulatory elements whose specific chromatin features remain to be identified.

## Background

Gene transcription is controlled by the combination of several classes of *cis*-regulatory elements, including promoters and enhancers. In addition to positive regulation, gene expression is controlled by negatively-acting regulatory elements known as transcriptional silencers. Silencers were first identified in *Saccharomyces cerevisiae* [1], and several decades later large-scale reporter assays identified large numbers of silencers in *Drosophila* [2], human cell lines [3,4], as well as cell lines and primary retina cells from mouse [5,6] and zebrafish [7].

While the prevalence and importance of silencers is now appreciated, the number of known silencers is still limited and their locations in the genome are difficult to predict. Enhancers can be predicted by the presence of specific histone modifications, most commonly H3K27ac [8] and H3K4me1 [9], although not all enhancers carry these marks. Others are marked by H3K9ac [10], H4K16ac [11], or the acetylation of lysines in the globular domain of histone 3, including lysines 64 and 122 [12]. Although no single mark predicts all enhancers, they can be predicted accurately based on the combination of these known marks together with peaks of chromatin accessibility and ChIP-seq for specific activator complexes [13]. In contrast, histone modifications indicative of silencers remain unknown. A known chromatin mark of silencers would enable researchers to predict this class of regulatory elements in any model system and cell type by a high-throughput sequencing assay of the characteristic chromatin mark (*e.g.*, ChIP-seq [14], CUT&Tag [15]).

Silencers active in *Drosophila* embryonic mesoderm were found to be significantly enriched for H3K27me3, but this modification was neither sensitive nor specific in distinguishing silencers: many silencers had low levels of mesodermal H3K27me3 and many non-silencers had high levels [2]. Ngan *et al.* identified silencers corresponding to chromatin loops bound by PRC2, the complex that deposits H3K27me3 [5]. Although H3K27me3 is found at many silencers, it is not highly predictive since many of them do not carry this modification and large expanses of the genome are covered in broad H3K27me3 domains, making it difficult to identify active silencers from within regions of Polycomb-repressed chromatin.

Large-scale screens for silencers (repressive ability of silencer elements or “ReSE”) in human cell lines reported that silencers are significantly enriched for methylated histone H4 lysine 20 (H4K20me) [4]. The input open chromatin screened by ReSE was obtained by formaldehyde-assisted isolation of regulatory elements (FAIRE) in K562 cells based on the assumption that silencers, like enhancers, are active *cis*-regulatory elements and therefore reside in open chromatin. The resulting FAIRE chromatin was then cloned into a reporter vector upstream of a toxic caspase 9 selection marker so that only cells with an active silencer survive screens performed in K562 and HepG2 cells. The authors then conducted an analysis for co-associated histone modifications using available ENCODE ChIP-seq peaks [16] and found that H4K20me was the most highly enriched mark in both cell lines [4], but the exact role of this modification remains unclear. We note that while their study refers to H4K20me generally and does not specify methyl form, the antibody used to generate the ENCODE ChIP-Seq data (Abcam ab9051) primarily detects monomethylation of H4K20me (H4K20me1)).

Other studies have found H4K20me1 to be associated with both repressed and actively transcribed chromatin. The mammalian inactive X chromosome is covered by H4K20me1 following binding of the long noncoding RNA Xist that silences the X chromosome, although H4K20me1 was not found to be essential for silencing of genes on the X chromosome [17].

H4K20me1 is also found on the inactive X chromosome of *C. elegans*, which uses an orthogonal dosage compensation mechanism whereby expression of each X chromosome in XX animals is reduced by half to match the expression of XO animals. In worms, H4K20me1 is enriched on both X chromosomes through demethylation of H4K20me2 by the demethylase DPY-21 [18]. While loss of DPY-21 catalytic activity led to derepression and decompaction of the inactive X, it was not lethal, again indicating H4K20me1 is not essential for viability.

H4K20me1 has been shown to directly recruit L3MBTL1 *in vitro* to promote chromatin compaction [19]. Nevertheless, the precise role of H4K20me1 in negative regulation of gene expression, particularly beyond X chromosome inactivation, is understudied. Conversely, H4K20me1 marks actively transcribed genes, along with H3K79me2 [20]. Beyond the transcribed gene body, H4K20me1 is commonly seen at promoters of expressed genes [21].

The connection between H4K20me1 and active transcription seems to run counter to the suggestion that this modification marks transcriptional silencers, but it is consistent with the finding that many silencers are bifunctional [2,5], acting as enhancers in alternate cellular contexts. One model from these data is that H4K20me1 marks bifunctional elements while they are active as both enhancers and silencers, leading to the observed bifunctional association of this mark. However, the most common form of H4K20 in the human and fly genomes is H4K20me2, comprising approximately 80% of histone tails, whereas H4K20me1 comprises approximately 10% of histone tails [22,23]. This presents technical difficulties for assaying H4K20me1 by ChIP-seq because even a small degree of cross-reactivity of antibody against H4K20me1 to the dimethyl form may greatly skew the results to H4K20me2, obscuring the signal due to H4K20me1.

Here, we report an in-depth re-analysis of histone mark enrichment among the human silencers identified by Pang and Snyder [4]. Whereas the authors of that study [4] calculated enrichment as compared to random sampling of the human genome, we instead used elements that were tested but not found to act as silencers. This method creates a more appropriate background for calculating enrichment since this maintains the sequence biases inherent in generating the FAIRE open chromatin library used to screen for silencers in that study. In contrast to the high degree of enrichment reported by Pang and Snyder [4], the results of our re-analysis indicate that H4K20me1 is only marginally enriched among the Pang & Snyder human silencers. We confirmed the generality of the lack of notable enrichment among silencers in *Drosophila* by generating H4K20me1 ChIP-seq data in *Drosophila* S2 cells using 3 different H4K20me1 antibodies. Despite extensive recent studies, silencers remain a poorly annotated class of *cis*-regulatory elements that cannot be reliably predicted by commonly assayed histone modifications.

## Results

### H4K20me1 is only marginally enriched among human K562 and HepG2 silencers

The authors of the ReSE silencer screen reported that H4K20me-modified chromatin was significantly enriched among silencers from K562 and HepG2 cells. We noticed that in their analyses, Pang & Snyder calculated the significance of enrichment for any histone modification using a permutation test in which silencer coordinates were randomly shuffled across the genome. However, since the FAIRE fragments used to assemble the tested library are enriched for open chromatin, the use of random genomic regions as background is not appropriate (Fig. 1A).

**Figure 1.**
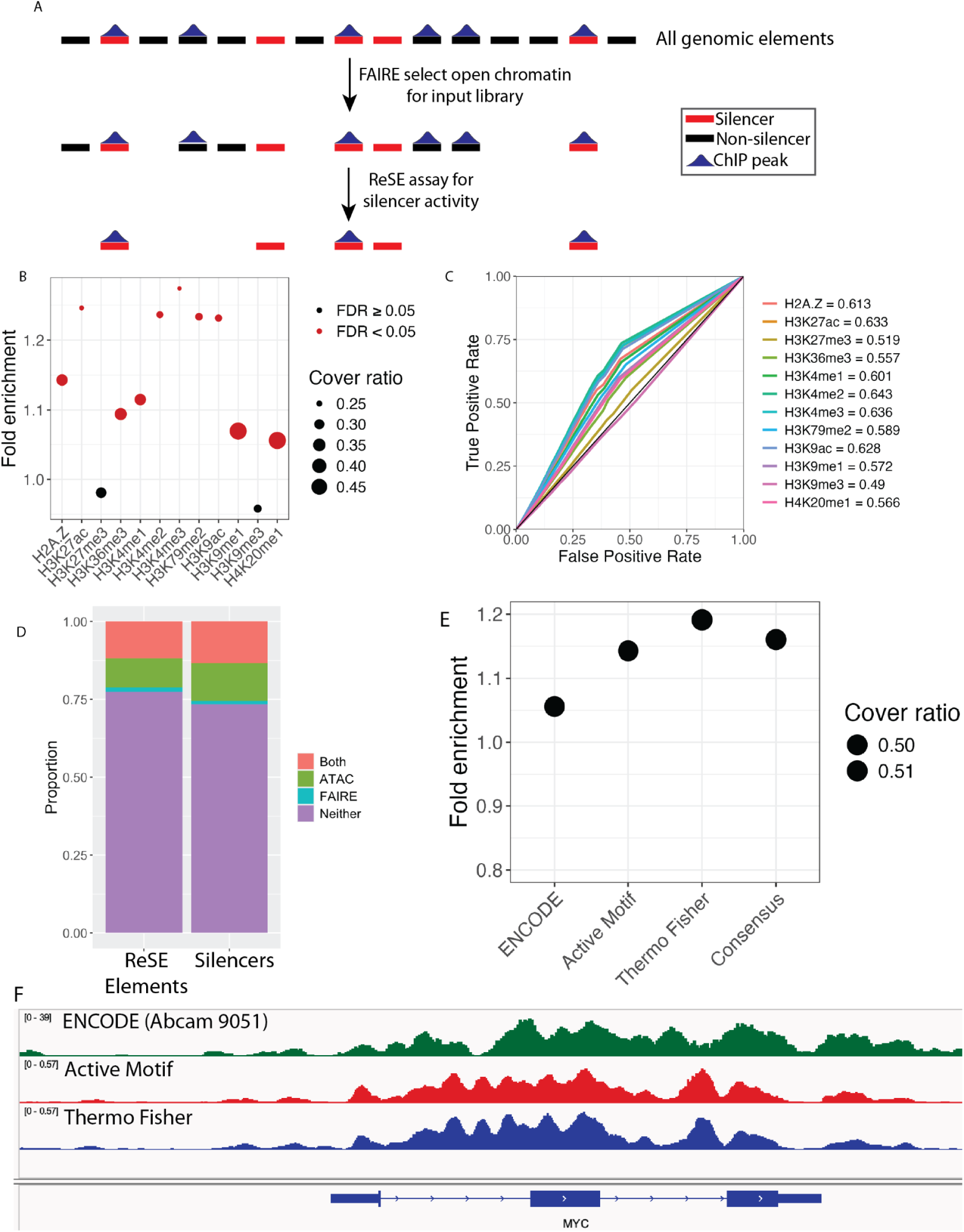
Silencers in human and fly cell lines are not enriched for H4K20me1. (**A**) Schematic of how FAIRE selection of an input library can create spurious enrichments. Top: all possible elements across the genome. Blue peaks represent histone modification ChIP-Seq peaks. Middle: selection of a subset of elements as selected by FAIRE for testing by ReSE enriches for histone modifications associated with open chromatin. Bottom: subset of tested elements that have significant silencer activity by MPRA. Proportion of elements overlapping the histone modification is unchanged compared to FAIRE elements but appears enriched compared to all possible genomic elements. (**B**) Results of permutation test enrichment analysis for the K562 silencers. The background elements were randomly selected from tested non-silencer elements. Cover ratio denotes the fraction of silencer elements overlapping the indicated ChIP-Seq peak set. Fold enrichment represents the cover ratio of the foreground silencers over the background cover ratio of non-silencer elements. *P*-values were computed empirically by a permutation test and adjusted by Benjamini-Hochberg correction to calculate a false discovery rate (FDR). (**C**) AUROC curves showing histone modification enrichments for K562 silencers. AUROC curves were computed by iterating over silencer score quantile cutoffs from the set of all tested elements in 1 percentile increments. (**D**) Proportion of all tested ReSE elements (left) and active silencers elements (right) that overlap accessible chromatin peaks called by FAIRE-seq (blue), ATAC-seq (green), or both (red). Regions that do not overlap accessibility peaks are purple. (**E**) Results of permutation test enrichment analysis as in **B** comparing results from H4K20me1 peak sets. (**F**) Genome browser tracks comparing ENCODE Consortium data using Abcam ab9051 (green) H4K20me1 with K562 ChIP-seq tracks using Active Motif AB_2615074 (red) or Thermo Fisher MA5-18067 (blue) antibodies. ENCODE track shows read depth, Active Motif and Thermo Fisher tracks are CPM scaled.

Therefore, we performed a similar permutation test except that to assemble a putative negative set we instead randomly sampled elements tested in the ReSE assay that were not found to exhibit silencer activity. The results revealed a strikingly different pattern of histone modification enrichment. In contrast to the results reported by Pang & Snyder [4], H4K20me1 was only moderately enriched (*q* < 0.005, 1.056-fold enrichment) and H3K27me3 and H3K9me3 were not enriched (*q* > 0.05, 0.958-fold enrichment). Instead, the strongest enrichments corresponded to activation-associated marks, including H3K27ac (1.246-fold enrichment, *q* = 0), H3K4me1/2/3 (1.115, 1.237, 1.74-fold enrichment, *q* = 0.00017, 0, 0, respectively) and H3K9ac (1.232-fold enrichment, *q* = 0) (Fig. 1B). However, despite the statistical significance, the magnitude of these enrichments was modest, with none enriched more than 1.25-fold over background. For comparison, a similarly constructed STARR-seq assay that selected open chromatin as input, but measured enhancer activity, resulted in at least 5 - 10 fold enrichment of enhancer-associated marks [24]. Although Pang & Snyder noted small but significant enrichment for H3K9me3, we instead observed slight but not statistically significant (*q* = 1.0 and 0.22, respectively) depletion of this mark among K562 and HepG2 silencers. Since ReSE employed a survival marker for detecting silencers, the assay was not able to differentiate between fragments with enhancer activity and fragments with no regulatory activity. This is likely why there are no strong depletions for marks commonly associated with enhancers. We observed very similar enrichments for silencers found by ReSE in HepG2 cells (Additional file 1: Fig. S1A); nevertheless, use of a FAIRE library from K562 cells as input for the ReSE screen in HepG2 cells likely biased the set of identified HepG2 silencers, and thus potentially also the associated histone marks.

Next, to evaluate the predictive accuracy of each of the histone modifications in distinguishing silencers versus non-silencers, we analyzed the K562 and HepG2 silencers by the area under receiver operating characteristic curve (AUROC) (see the “Methods” section), which assessed performance across all possible threshold scores for calling silencers from the ReSE data. For the background in this AUROC analysis, as above, we used randomly sampled elements tested in the ReSE assay that were not found to exhibit silencer activity. The AUROC results were overall consistent with those from our permutation tests: the strongest, albeit modest, enrichments were for enhancer-associated marks, including H3K4me1/2/3 and H3K9ac (AUROC = 0.601, 0.643, 0.636 and 0.628, respectively; Fig. 2C), while H3K9me3 was slightly depleted (AUROC = 0.490). We obtained similar results for silencers found by ReSE in HepG2 cells, with an even more pronounced depletion of H3K9me3 among the HepG2 silencers (AUROC = 0.417; Additional file 1: Fig. S1B).

**Figure 2.**
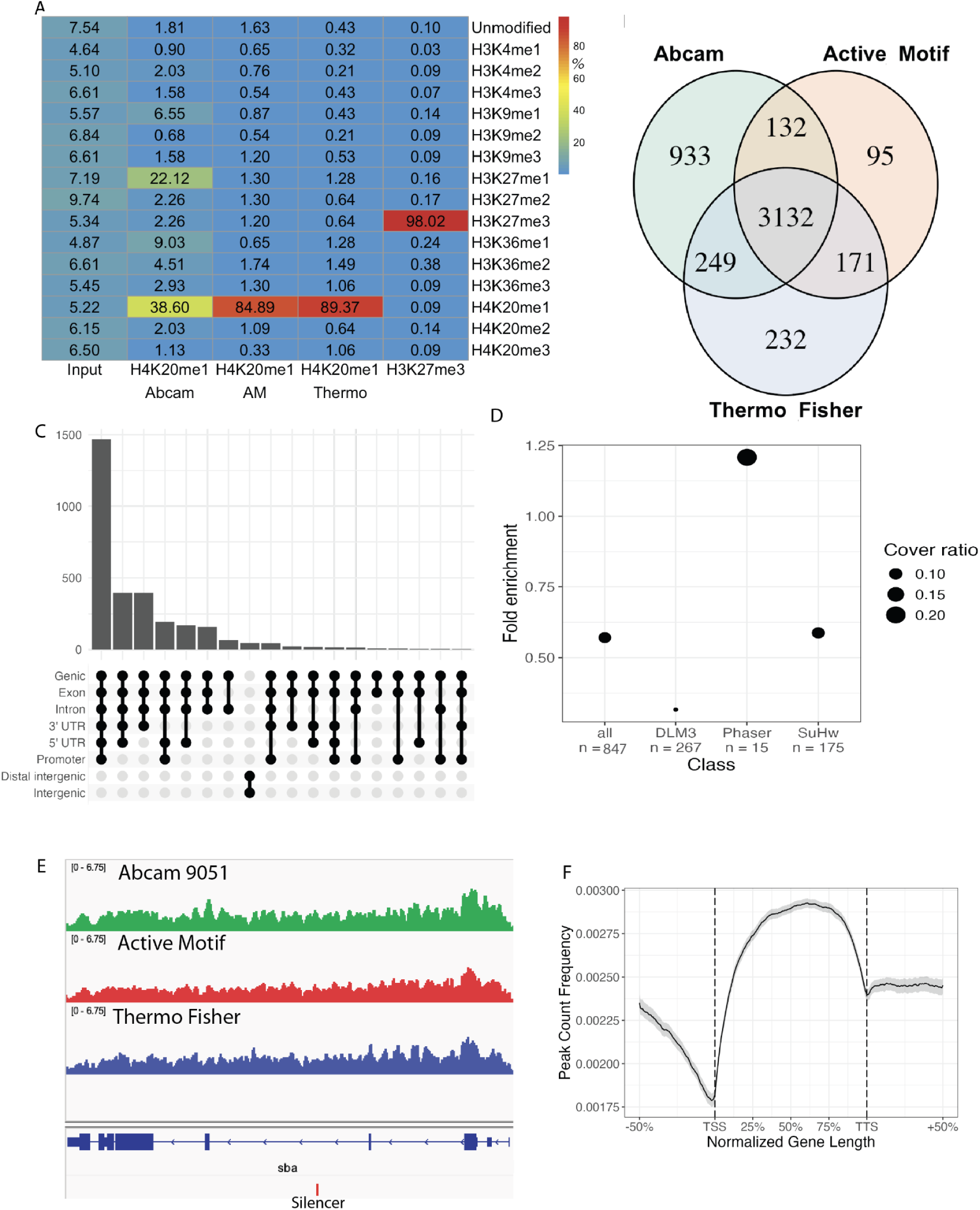
H4K20me1 marks actively transcribed genes in *Drosophila* S2 cells. **(A)** Summary of antibody specificity as measured by SNAP-ChIP spike-in mononucleosomes. Columns represent individual ChIP-seq libraries, rows represent all histone modifications present in the panel. Values denote percentages of barcode reads mapped to each modification within a library. Abcam 9051: Abcam ab9051; AM: Active Motif AB_2615074; Thermo Fisher: Thermo Fisher MA5-18067. (**B)**. Venn diagram comparing IDR reproducible peaks in S2 cells from the three H4K20me1 antibodies used in this study. (**C)** Upset plot of genomic annotations for *Drosophila* S2 cell H4K20me1 ChIP-Seq peaks using Ensembl gene annotations for dm3 (**D)** Results of H4K20me1 permutation test enrichment for *Drosophila* S2 silencers. DLM3, Phaser, and Su(Hw) are subsets of “all” silencers that contain matches for the named motif (*e.g.*, DLM3) and are not mutually exclusive. *p*-values were computed by permutation tests using random genomic regions. None of the classes were significantly enriched for H4K20me1 at *P* < 0.1. All: n = 837; DLM3: n = 267; SuHw: n = 175; Phaser: n = 15. Average peak density of H4K20me1 consensus ChIP-Seq peaks over genes. (**E)** Genome browser tracks CPM normalized read depth for H4K20me1 ChIP-seq in S2 cells using Abcam ab9051 (green), Active Motif AB_2615074 (red), and Thermo Fisher MA5-18067 (blue) antibodies. The red tick below the gene model shows the location of a silencer marked by H4K20me1. (**F**) Aggregate peak profile for H4K20me1 consensus peaks in S2 cells over genes annotated in Ensembl for dm3.

Because the input library was obtained by FAIRE, we reasoned that FAIRE data from ENCODE for K562 cells [16] might serve as an alternate background set (without the more stringent selection of a putative negative set) in these histone mark enrichment analyses. The resulting profile of histone modifications enriched among the K562 silencers was vastly different using the ENCODE FAIRE sequences as background rather than the putative non-silencers from the ReSE screen. Here, only H3K27me3 was significantly enriched (*q* < 0.05), again with a modest enrichment (Additional file 1: Fig. S2A). Only 13% of ReSE input fragments overlapped with ENCODE FAIRE peaks from K562 cells, while many more sequences were present only in the ReSE input library or the ENCODE FAIRE data (Fig. 1D). The ReSE input library showed lower overlap with HepG2 FAIRE-seq peaks (7.5%), as expected since the ReSE library was prepared from FAIRE sequences obtained from K562 cells. Similarly, analysis of the HepG2 silencers using the HepG2 FAIRE-seq peaks as background showed significant enrichments only for H3K27me3 and H3K9me3 (Additional file 1: Fig. S2B). These results suggest that enrichment analyses of ReSE data should be performed for background sequences obtained from the same FAIRE experiments used to generate the ReSE input library to control for systematic biases across FAIRE datasets. Both the ReSE input library and the ENCODE FAIRE data showed only modest overlap with ENCODE ATAC-seq, despite both FAIRE and ATAC-seq targeting open chromatin (Fig. 1D), consistent with prior studies that reported differences in open chromatin regions obtained by different chromatin accessibility profiling technologies [25]. Conservatively considering only ReSE library elements that overlapped with FAIRE-seq peaks to assemble a background set, enrichments for histone modifications showed even more modest effect sizes (fold enrichment < 1.05, *q* > 0.05 for all marks, Additional file 1: Fig. S2C). Similarly, only 11% of silencers identified in HepG2 overlapped with ENCODE FAIRE peaks; restricting analyses to these regions did not substantially change the enrichment results (Additional file 1: Fig. S2D).

Since the histone modifications that we found to be most enriched among the K562 or HepG2 silencers – H3K4me1/2/3 and H3K9ac – are marks associated with promoters [8,26], we reasoned that this enrichment could be driven by promoter competition [27] and thus excluded promoter regions from further analyses of silencer features [2]. Inspection of the distributions of the top 1,000 ENCODE ChIP-Seq peaks for the promoter-associated marks (H3K4me2/3 and H3K9ac) showed that these peaks are generally located within 5 kb of transcription start sites (TSSs) (Additional file 1: Fig. S3A-C). Therefore, to focus our analysis of potential silencer marks beyond those whose silencing activity is likely due to promoter competition, we omitted all ReSE library elements within 5 kb of a TSS. Although both the statistical significance and fold enrichment for H3K4me2/3 were diminished, the degree of those reductions was small and H3K4me2/3 remained among the most highly enriched marks among the K562 silencers (Additional file 1: Fig. S3D-E).

### *Drosophila* S2 cell silencers are not enriched for H4K20me1

To validate the findings from human cell lines in an orthologous system, we next looked to silencers in *Drosophila* melanogaster. To date, the largest set of silencers identified in *Drosophila* was found using a STARR-seq-based screen in S2 cells [28]. This screen identified 837 active silencer elements, many of which contain known DNA binding site motifs for the transcriptional repressors Phaser or Su(Hw) or the uncharacterized motif DLM3 [29].

Because H4K20me1 ChIP-seq data for S2 cells were not publicly available, we generated ChIP-seq data in S2 cells using three antibodies against H4K20me1 and one antibody against H3K27me3 as a positive control (see the “Methods” section). The H3K27me3 ChIP-seq peaks had strong overlap with previously published H3K27me3 ChIP-seq data for S2 cells [30] (Additional file 1: Fig. S4), which suggests the ChIP-seq experiments worked successfully. To directly assay the specificity of each antibody, we utilized SNAP-ChIP spike-in mononucleosomes, a panel of modified mononucleosomes with DNA barcodes representing specific modifications [31]. SNAP-ChIP spike-in mononucleosomes showed that all three H4K20me1 antibodies enriched H4K20me1 nucleosomes with low cross-reactivity (∼0.6 to 3.5%) with the di- or tri-methylated forms (H4K20me2/3), although one of the antibodies (Abcam 9051) exhibited substantial cross-reactivity (>55%) with other monomethylated lysines, particularly H3K27me1 (Fig. 2A). This cross-reactivity suggests that many peaks in prior ChIP-seq datasets generated using this antibody, including the ENCODE ChIP-seq data for K562 and HepG2 [16] used to analyze the ReSE silencer data by Pang and Snyder [4] and by us in this study, may not actually be due to H4K20me1.

IDR analysis showed good reproducibility (36.7% - 50.1% of peaks replicated at IDR < 0.05, Additional file 1: Table S2) between replicate ChIP-Seq experiments (Additional file 1: Fig. S5). Furthermore, the sets of IDR peaks resulting from each of the three antibodies were largely overlapping (Fig. 2B). Abcam 9051 had the largest number of peaks not found using the other antibodies, most likely because of the high cross-reactivity of this antibody. We merged the peaks found in common by all three H4K20me1 antibodies into a high-confidence consensus set of 3036 H4K20me1 peaks (see the “Methods” section) with an average peak width of 9 kb. 1486/3036 (49%) of these H4K20me1 consensus peaks overlapped peaks identified by ChIP-chip experiments by modENCODE [32] (Additional file 1: Fig. S6A). This moderate degree of overlap may be due to technical differences between the ChIP-Seq and ChIP-chip experiments and associated peak calling algorithms. The strongest modENCODE ChIP-chip peaks showed the greatest ChIP-Seq peak signal (CPM) from the overlapping H4K20me1 consensus ChIP-Seq peaks (Additional file 1: Fig. S6B), suggesting that the weaker modENCODE peaks may have been more likely to be due to antibody cross-reactivity. H4K20me1 peaks covered a smaller proportion of the *Drosophila* genome overall (∼17% of mappable dm3 genome) compared to K562 (∼28% of mappable hg19 genome). This is most likely the result of our more stringently requiring H4K20me1 peaks in S2 cells to have been found using three different antibodies, but might also reflect broad, species-specific differences.

Using our high-confidence consensus set of H4K20me1 ChIP-Seq peaks from S2 cells, we analyzed the set of 837 S2 silencers for enrichment of H4K20me1 ChIP-Seq peaks. In contrast to the FAIRE-enriched library used as input in ReSE, the S2 silencer screen was constructed using random fragments covering the entire *Drosophila* genome; therefore, we used random *D. melanogaster* genomic sequences as a proxy for an input library [28]. We found that the S2 silencers as a whole were not enriched for H4K20me1 peaks (Fig. 2D). We next analyzed each of the 3 classes of silencers, as defined according to their repressor motif matches (DLM3, Phaser, Su(Hw)), for enrichment of H4K20me1 to test if this histone modification marks a specific class of silencers. None of these 3 silencer classes were significantly enriched for H4K20me1 (*P* < 0.05 after Benjamini-Hochberg correction to adjust for multiple hypothesis testing). Though not statistically significant, silencers with Phaser motifs showed an enrichment of 1.21-fold (Figure 2D). However, this subclass had a very small number of silencers (n = 15), only 3 of which were marked by H4K20me1 peaks, and all of these peaks were within introns of expressed genes (Figure 2E), which suggests that H4K20me1 marked transcription in these instances rather than marking silencer activity. As an alternative approach for calculating enrichment, we generated a background set matched for genomic sequence composition using GENRE [33]. We found that silencers had a comparable degree of enrichment for H4K20me1 using GENRE background compared to random background (Additional file 1: Fig. S7). Notably, the cover ratio of H4K20me1 over S2 silencers (0.09) was lower than that observed in human K562 and HepG2 cells (0.50 and 0.40 respectively).

### H4K20me1 is associated with active transcription in S2 cells

Genomic annotations of H4K20me1 peaks in S2 cells showed these regions are primarily genic and cover both introns and exons (Fig. 2C). Peaks were generally within 1 kb of a TSS (Additional file 1: Fig. S8). Only 4 peaks were annotated as “distal intergenic”, and inspection of these loci showed that they often included lncRNAs that were not included in the Flybase gene annotation file but are presumably transcribed. ENCODE H4K20me1 peaks in a human cell line (K562) showed broadly similar distributions across different genomic regions when considering the much larger genome and longer intronic spans of human genes (Additional file 1: Fig. S9). In both *Drosophila* S2 cells and human K562 cells, the profile of H4K20me1 over gene bodies shows a dip directly over TSSs before sharply increasing and slowly attenuating over the length of the gene body, consistent with a prior study of H4K20me1 in human cell lines (Fig. 2F and Additional file 1: Fig. S9) [20].

Since H4K20me1 previously was found to be enriched over transcribed genes in mammalian cells and *Drosophila* [20,34–36], we integrated RNA-seq expression data from S2 cells into our analysis to inspect H4K20me1 for potential association with gene expression levels [37]. We found that genes overlapping S2 cell H4K20me1 ChIP-Seq peaks were on average more highly expressed in S2 cells than were genes that did not overlap H4K20me1 ChIP-Seq peaks (*P* < 2.2 x 10^-16^) (Additional file 1: Fig. S10). Analysis of Gene Ontology (GO) annotation terms assigned to H4K20me1 intersecting genes showed enrichment for developmental categories, including “instar larval or pupal morphogenesis”, “imaginal disc morphogenesis” and “wing disc development” (Additional file 1: Fig. S11), consistent with the stem cell origins of S2 cells [38]. Several GO categories related to negative regulation were enriched, including “negative regulation of response to stimulus”, “negative regulation of signaling” and “negative regulation of cell communication”, suggesting that many H4K20me1 marked genes are involved in regulating cellular responses. Profiles of H4K20me1 have been seen to closely match those of H3K36me3, another mark of actively transcribed gene bodies, in human cell lines [20]. We analyzed publicly available H3K36me3 ChIP-seq peaks from S2 cells [30] and found that 60% of H4K20me1 ChIP-Seq peaks overlapped with H3K36me3 peaks (Additional file 1: Fig. S12A), demonstrating the evolutionary conservation of the co-association of these marks with expressed genes. The distribution of these H3K36me3 peaks was strikingly similar to that of H4K20me1 in terms of distance to TSS, genomic annotation, and aggregate peak profile over gene bodies (Additional file 1: Fig. S12B-D). Overall, we found that the distribution and associations of H4K20me1 in S2 cells is similar to known H4K20me1 patterns in other systems.

### Degree of H4K20me1 enrichment among silencers is not dependent on antibody choice

Because the Abcam H4K20me1 antibody showed a high degree of cross-reactivity in *Drosophila* cell ChIP-seq libraries (Fig. 2A), we next investigated whether the lack of enrichment among human silencers might be due to low signal to noise in the ENCODE reference dataset. We performed H4K20me1 ChIP-seq experiments for K562 cells using the two H4K20me1 antibodies that showed high specificity for H4K20me1, namely those from Active Motif and Thermo Fisher, hereafter referred to as “Thermo” for simplicity in writing (Methods). SNAP-ChIP modified nucleosome spike-ins showed that both antibodies (93.39% for Active Motif and 78.33% for Thermo) (Additional File 1: Fig. S13A) had high on-target specificity. Replicate ChIP-seq data for both antibodies showed very high correlation of ChIP-seq signal across all genomic bins (Spearman ρ = 0.98 - 0.99), with the ENCODE H4K20me1 track showing a consistent, moderate pairwise correlation with all the new ChIP-Seq data (Spearman ρ = 0.77 - 0.81) (Additional File 1: Fig. S13B).

Peak calling with MACS2 revealed very similar numbers of peaks for both new antibodies (264,921 and 268,677 IDR reproducible peaks for Active Motif and Thermo, respectively) (Additional File 1: Table S4). 64% of ENCODE peaks were reproduced by both Active Motif and Thermo antibodies (Additional File 1: Fig. S13C), suggesting that some ENCODE peaks may be attributable to antibody cross-reactivity. We generated a consensus set of H4K20me1 peaks that are present in both Active Motif and Thermo peak sets, for a total of 190,627 high-confidence peaks. Similar to previous H4K20me1 datasets, the H4K20me1 consensus peaks were found primarily in genic regions and near TSSs (Additional File 1: Fig. S13D-E). The distribution of H4K20me1 signal over gene bodies showed the characteristic profile, with broad peak that gradually declines over the length of the gene body (Fig. 1F and Additional File 1: Fig. S13F). Similar to the S2 H4K20me1 dataset, H4K20me1 peaks in K562 cells showed substantial overlap with H3K36me3 peaks (26% of H4K20me1 peaks) (Additional File 1: Fig. S14A), with H3K36me3 signal showing a similar genic distribution and profile over genes (Additional File 1: Fig. S14C-D), consistent with H4K20me1’s association with active transcription. We found very similar results from permutation tests for enrichment of H4K20me1 among silencers when using ENCODE, Active Motif, Thermo, or consensus peak sets (Fig. 1E), indicating that our observation that H4K20me1 is only marginally enriched among human K562 silencers, is robust and not substantially affected by antibody cross-reactivity.

## Discussion

Human transcriptional silencers were reported previously to be enriched for H4K20me1 [4], but our re-analyses of those human silencer data using appropriate background sets to assess enrichment, combined with analysis of *Drosophila* S2 cell silencer data with H4K20me1 ChIP-Seq profiles generated for this study, reveal that H4K20me1 is not actually a silencer mark in either human or fly. Our analyses demonstrate the importance of both selecting the appropriate background in enrichment analyses and considering the fold enrichment to evaluate the effect size, in addition to statistical significance. Although the enrichment of some histone modifications did reach statistical significance, their modest fold enrichments indicate that they are not reliable predictors. Though not highly enriched, the presence of marks typically associated with enhancers, such as H3K27ac and H3K4me2, on silencers was surprising. Our results are consistent with a model whereby a subset of silencers with these marks are associated with active nuclear compartments containing target genes. In S2 cells, however, these activating marks were found to be depleted among silencers [28], suggesting that this model may be species-specific or alternatively applies to only a small subset of silencers.

The ReSE library obtained by FAIRE in K562 cells [4], which was used as input in the human silencer screen that we re-analyzed, showed surprisingly low overlap with ENCODE FAIRE-seq peaks from the same cell line (Fig. 1D). However, only a small portion of reads (FAIRE fragments) in a FAIRE-seq experiment are typically found within peaks [39], consistent with the fraction of FAIRE fragments from ReSE that were in FAIRE-seq peaks. ReSE [4], as well as several other silencer screens [2,3,40], selected for open chromatin based on the assumption that silencers are located within accessible chromatin, as are transcriptional enhancers. The result that most K562 and HepG2 silencers were not found within open chromatin according to FAIRE-Seq or ATAC-seq data challenges this assumption and suggests that human silencers either do not reside preferentially in open chromatin or alternatively that they require only very short stretches of open chromatin that may not be detected by genomic assays such as FAIRE-seq. Such a chromatin state is consistent with findings from the recent *Drosophila* S2 cell silencer screen, which found that that *Drosophila* S2 silencers reside within phased nucleosomes that are present within inaccessible chromatin, according to DNase I hypersensitivity peaks, but often contained very short regions of chromatin accessibility that were sufficient for just a single transcription factor to bind [28].

Together, these findings suggest that histone modifications may not be a reliable predictor of silencers and that new strategies may be needed to identify silencers. Although only a subset of silencers are marked by H3K27me3, many silencers have been found by mapping chromatin loops anchored by H3K27me3 or PRC2 [5,41]. Thus, H3K27me3 Hi-ChIP could be used to identify additional silencers, albeit only for a subset of silencers. A general chromatin feature of silencers that has been identified thus far is the binding of a specific transcriptional repressor within a narrow region of chromatin accessibility [28,41,42]. Under this model, silencers active in a cell type of interest would be predicted by intersecting high-resolution chromatin accessibility data (*e.g.*, DNase-seq) displaying such an accessibility profile in that cell type with motif matches for known repressors expressed in that cell type. Massively parallel reporter assays, particularly those that screen the entire genome, remain the most comprehensive experiments to identify novel silencers. The S2 silencer screen [28] highlighted the power of coupling DNA affinity purification with mass spectrometry by identifying a novel repressor essential for the activity of a large fraction of S2 silencers. When coupled with locus-specific purification [42,43] or proximity labeling [44], mass spectrometry would offer a way to identify factors involved in silencing and associated histone modifications. Alternatively, a CRISPR screen could be used to identify genes necessary to maintain repression of a reporter under the control of a silencer; the resulting candidate silencer regulators, if found to interact physically with silencers, could be tested for their utility in predicting silencers. Highlighting their high degree of bifunctionality, many silencers have been found among annotated enhancers from other cell types [2,5,41], suggesting annotation as an enhancer in another cell type as a predictive feature of silencers [2]. Future studies are needed to determine if there is a combination of various genomic features (*e.g.*, chromatin accessibility, histone modifications, motif composition, sequence annotation, nascent transcription) that could be used to predict a wide range of silencers.

Our results highlight persistent problems with antibody specificity. Poor antibody specificity can lead to reduced signal to noise ratio or false positive ChIP-seq peak calls. While some of the antibodies used here were highly specific, the widely used H4K20me1 antibody Abcam 9051 showed substantial off-target affinity for other monomethylated lysines, especially H3K27me1. This cross-reactivity could create confounding artifacts in interpreting results from ChIP-seq and other experiments using this antibody, since H3K27me1 is related to the facultative heterochromatin mark H3K27me3 and is also deposited by PRC2 [45]. This antibody has been discontinued but was widely used in many previous studies, including by ENCODE and modENCODE. Because ChIP-seq studies rarely include direct assays for antibody specificity, such as the barcoded mononucleosomes used here, antibody cross-reactivity often goes unnoticed. To address this issue, we have provided new ChIP-seq data for H4K20me1 in K562 cells with more specific antibodies that we validated by spike-in mononucleosomes. While the main results of our study were not substantially affected by antibody choice, these new H4K20me1 ChIP-seq data will be a valuable resource for future studies investigating the functional associations of this mark.

The functional role(s) of H4K20me1 remains unclear. The loss of H4K20me1 results in reduced viability [33] or reduced efficiency of X chromosome inactivation [13], suggesting that H4K20me1 may contribute to robustness of gene regulatory programs. While it has been found to localize on regions of heterochromatin [17,19,46], our results from H4K20me1 ChIP-seq in S2 cells support prior studies in human cells that found it associated with actively transcribed genes [20,35,47]. Our data showed association almost exclusively with active chromatin, but we note that many of the previous associations with repressed chromatin, such as the mammalian inactive X chromosome [17] or polytene chromosomes [46], are not present in S2 cells. H4K20me1 is also associated with the repression of repetitive elements in mammals [48,49], but *Drosophila* primarily use small RNA pathways to silence repetitive elements [50], for which we did not find an association with H4K20me1.

## Conclusions

Our results expand upon a recent study in *Drosophila* S2 cells that likewise found that transcriptional silencers do not correspond to any known chromatin signatures [28]. Despite the number of functionally identified silencers in fly or human now numbering in the thousands [3–5,28], they remain poorly characterized and difficult to predict. Silencers might be specifically marked by a histone modification that is not commonly profiled or for which ChIP-grade antibodies do not exist currently. Mass spectrometry has identified over 500 histone post-translational modifications [51], only a small fraction of which have been assayed by ChIP-seq or similar assays. The functional associations of many of these understudied modifications, particularly those other than methyl or acetyl groups, have started to be appreciated only recently [52,53]. Broader profiling of histone modifications may reveal a silencer chromatin signature. Several distinct subclasses of silencers may exist, each of which are characterized by a different chromatin signature such that there is no universal silencer mark [54]. Finally, it is possible that unlike transcriptional enhancers, silencers as a broad class of *cis*-regulatory elements have no characteristic histone mark(s) and can only be predicted by the binding of particular transcriptional (co-)repressors. It also remains unclear the extent to which the chromatin state of silencers is dynamic or whether it is affected by cell cycle or other aspects of cellular context. Determining the chromatin features of silencers will be important for the prediction of transcriptional silencers for genome annotation and understanding gene regulatory mechanisms.

## Methods

### S2 cell culture

S2-DRSC cells were purchased from the *Drosophila* Genomics Resource Center (DGRC Stock Number: 181). S2 cells were cultured in Schneider’s medium (Thermo Fisher 21720024) supplemented with 10% fetal bovine serum (FBS) and penicillin-streptomycin at ambient temperature (22 °C). Cell counts were determined using a Countess II with Trypan blue staining.

### ChIP-seq

We performed ChIP-seq experiments in S2 cells closely following previously published protocols [55]. Briefly, we used 100 µg of sheared DNA per ChIP reaction with SNAP-ChIP K-MetStat spike-in mononucleosomes. We used the following H4K20me1 antibodies: Abcam ab9051, used widely by ENCODE and modENCODE as well as [36]; Active Motif AB_2615074, which was not used in a published ChIP-seq study to our knowledge; and Thermo Fisher MA5-18067, which was used recently in CUT&RUN experiments in *Drosophila* [56]. As a positive control, we performed ChIP in parallel for H3K27me3 using a well characterized H3K27me3 antibody (Epicypher 13-0055). ChIP-seq reads were mapped to dm3 using Bowtie2 [57] and peaks were called using MACS2 [58]. We used the IDR framework [59] to find peaks that are reproducible between replicates.

### SNAP-ChIP spike-in analysis

We identified SNAP-ChIP barcodes in each library using a custom Python script based on Epicypher’s provided script. Counts were summed across the two redundant barcodes used for each modification, the two FASTQ files for each paired-end read library, and the two replicate libraries for each antibody.

### Statistical analysis

#### Enrichment of histone marks among genomic regions

For re-analysis of human datasets [4], enrichments were calculated using ChIP-seq data generated by ENCODE. Unless otherwise stated, all coordinates are in hg19. To match reference data used previously [4], processed bed files for broad peaks were downloaded from the UCSC Genome Browser. Reference peaks for H3K27me3 and H3K36me3 in S2 cells were downloaded from GEO: GSE245077 [30] and lifted over to dm3 coordinates. RNA-seq expression data for S2 cells were downloaded from SRA PRJNA937779 [37].

### AUROC analysis of histone mark enrichment among silencers

Silencer reporter activities for all tested human elements in ReSE were obtained from the authors of [4]. We performed receiver operating characteristic curve (AUROC) analysis to determine the sensitivity and specificity with which the silencer activity of an element is predicted by overlap (at least 1 bp) with a ChIP-seq peak for a particular histone modification. The entire tested library was sorted by fold enrichment and the cutoff values for every percentile were calculated. Then, for every percentile threshold, elements with fold enrichment greater than the threshold were considered silencers and those below the threshold considered non-silencers. The true positive (TP) elements were counted as those library elements above the silencer threshold and overlapping a ChIP-seq peak, false positives (FP) were above the silencer threshold but not overlapping a peak. True negatives (TN) were below the silencer threshold and not overlapping a peak, and false negatives (FN) were below threshold but overlapping a ChIP peak. The true positive rate (TPR) is TP/(TP+FN) and the false positive rate (FPR) is FP/(TN+FP). We computed the area under the AUROC (*i.e.*, the area under the curve of TPR as a function of FPR) by the AUC function of DescTools using the “trapezoid” method.

### Analysis of cover ratios of histone marks among human silencers

Cover ratio was defined as the fraction of foreground elements that overlap ChIP-Seq peaks [4]. We defined the fold enrichment as the ratio of the foreground cover ratio over the mean cover ratio of permuted background sets. Permutation tests were performed by comparing the fraction of elements overlapping annotated peaks in foreground (silencer) versus background (non-silencer) sets. For histone modification enrichment analyses in ReSE datasets, we defined the background as the set of elements tested in the screen that were not called as silencers.

Because the *Drosophila* S2 STARR-seq screen had much broader coverage across the genome in the input library, we defined the background as the entire *D. melanogaster* genome, excluding unmappable blacklisted regions. *P*-values of histone mark enrichment were computed as the fraction of permuted background sets that had a higher cover ratio than the foreground set. All permutation tests were performed using 20,000 permutations to mirror the analyses performed by Pang and Snyder [4].

### Analysis of S2 cell RNA-seq expression data

We downloaded a table of TPM normalized read counts from an S2 RNA-seq experiment [37]. We classified genes as H4K20me1+ if they intersected an H4K20me1 consensus peak by at least 1 bp and H4K20me1- if they did not. The statistical significance of differences between the distributions of TPMs among H4K20me1+ versus H4K20me1- genes was determined by a Student’s t-test.

### K562 cell line and culture

K562 cells were obtained through a Material Transfer Agreement (MTA) from the American Type Culture Collection (ATCC). Cells were cultured in RPMI 1640 (Corning 45000-396) supplemented with 10% Fetal Bovine Serum (FBS) and 1% Penicillin-Streptomycin-Glutamine (Thermo Scientific 10378016) at 37 °C with 5% CO_2_.

### ChIP-seq in K562 cells

#### Chromatin Isolation and Sonication

Wild type K562 cells were fixed for 5 minutes in Dubecco’s Phosphate Buffered Saline (1X DPBS, without Calcium and Magnesium, Corning 21-031-CV): complete media at a 2:1 ratio with addition of 1% formaldehyde. Crosslinking was quenched by the addition of glycine to a final concentration of 0.125 M, followed by a 5-minute incubation. Crosslinking and quenching reactions were both performed at room temperature with gentle agitation. Cells were collected on ice and washed with ice-cold 1X PBS before being resuspended in Sonication Buffer (20 mM Tris, pH 8.0, 2 mM EDTA, 0.5 mM EGTA, 1X Halt Protease and Phosphatase Inhibitor Cocktails (Thermo Scientific 78442), 0.5% SDS, and 0.5 mM Phenylmethylsulfonyl fluoride (PMSF)) at a concentration of 1 x 10^8^ cells per mL. Chromatin was sheared to an average size of ∼200 bp using a QSonica Q800R3 sonicator, flash-frozen in liquid nitrogen, and stored at -80 °C prior to immunoprecipitation.

#### Immunoprecipitation

ChIP samples were normalized by spiking with 2 µl SNAP-ChIP K-MetStat spike (Epicypher 19-1001) per 10 µg chromatin. Spiked material was diluted ∼1:10 in IP buffer (20 mM Tris pH 8.0, 2 mM EDTA, 0.5% Triton X-100, 150 mM NaCl, 10% Glycerol) and pre-cleared with 30 µL of Protein G (H4K20me1, Millipore 16-266) or protein A (H3K27me3, Millipore 16-125) agarose beads for ≥1 h at 4 °C. Pre-cleared samples were incubated overnight at 4 °C with 10 µg primary antibody per IP (H4K20me1: Thermo 5E10-D8 & Active Motif 39027; H3K27me3: EpiCypher 2084-1G5). Subsequently, 200 µL of pre-washed protein A/G bead slurry was added to each IP reaction and samples were incubated for ≥2 h at 4 °C. The post-IP washes, elution and reverse-crosslinking were performed as described in Martin *et al.* [60]. Samples were treated with Proteinase K (Thermo Scientific 25530049), extracted with phenol: chloroform: isoamyl alcohol (25:24:1 v/v, Invitrogen 15593049) and resuspended in 30 µL H2O following DNA precipitation.

#### Library Preparation

Libraries were prepared using the NEBNext Ultra II DNA Library Prep Kit for Illumina (New England Biolabs) according to the manufacturer’s instructions. Libraries were sequenced on an Illumina NovaSeq X Plus using a NovaSeq X Plus 10B lane with paired-end 151-bp cycles at the Biopolymers Facility at Harvard Medical School.

### Data processing

ChIP-seq reads were mapped to hg19 using Bowtie2 [57] and peaks were called using MACS2 [58]. We used the IDR framework [59] to find peaks that are reproducible between replicates.

Bigwigs were generated from deduplicated BAM files using Deeptools bamCoverage [61] with CPM normalization. Bigwigs for replicate samples were combined using Deeptools bigwigAverage.

## Supporting information

Table S5, Supplemental methods, Supplementary references

## Additional method details

For additional methods, please see Additional file 2: Supplemental Methods.

## Declarations

### Ethics approval and consent to participate

Not applicable.

### Consent for publication

Not applicable.

Availability of data and materials

Accession numbers of histone modification ChIP-Seq datasets for human cell lines from the UCSC Genome Browser analyzed in this study are listed in Table S3. Histone modification ChIP-Seq and RNA-Seq datasets for *Drosophila* S2 cells analyzed in this study were obtained from the NCBI Gene Expression Omnibus (GEO; http://www.ncbi.nlm.nih.gov/geo/) under accession number GSE245077 and the Sequence Read Archive (SRA) under accession number PRJNA937779, respectively. Genomic coordinates of S2 silencers were obtained from [28] under GEO accession number GSE254776. The raw sequencing data from H4K20me1 ChIP-Seq for S2 and K562 cells generated in this study have been deposited in GEO under accession number GSE284058. The results of all statistical tests appear in Additional file 3: Table S6-17. Original code for enrichment tests is available in Additional file 4.

### Funding

This project was supported in part by fellowship F31 GM145107 from the U.S. National Institutes of Health to J.A.S., grants R01 HG009723 and R56 HG009723 from the U.S. National Institutes of Health to M.L.B., grant R01 GM134539 from the U.S. National Institutes of Health to K.A., and a grant from the Brigham Research Institute Fund to Sustain Research Excellence to M.L.B.

Authors’ contributions

J.A.S. and M.L.B. designed research; J.A.S. and N.A.M performed experiments; J.A.S. performed data analysis; M.L.B. and K.A. supervised research; J.A.S. and M.L.B. wrote the paper. The authors read and approved the final manuscript.

## Acknowledgements

We thank Steve Gisselbrecht and HyuckJoon Kang for technical assistance, Steve Gisselbrecht and Luca Mariani for assistance with computational analyses, Kaia Mattioli for helpful discussion, and Steve Gisselbrecht, Kaia Mattioli and Julia Rogers for critical reading of the manuscript. We thank Alex Stark for very kindly sharing the genomic coordinates of the 837 *Drosophila* S2 cell silencers identified in their STARR-Seq-based screen prior to the publication of their paper presenting those silencers.

