## Supplementary material for "Histone H4 lysine 20 monomethylation is not a mark of transcriptional silencers": Figs S1-S14, Tables S1-S4

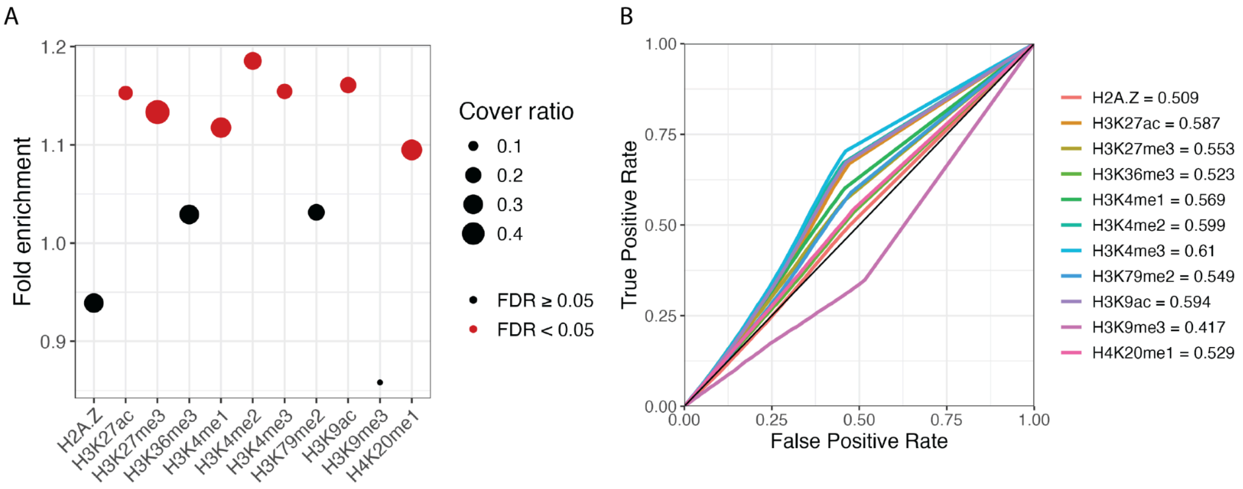


**Figure S1: HepG2 silencers are not enriched for H4K20me1.** (**A**) The background elements were randomly selected from tested non-silencer elements. Cover ratio denotes the fraction of silencer elements overlapping the indicated peak set. Fold enrichment represents the cover ratio of the foreground silencers over the background cover ratio of non-silencer elements. *P*-values were computed empirically by a permutation test with 20,000 permutations of the background set and FDR corrected by Benjamini-Hochberg. (**B**) AUROC-based enrichment scores for HepG2 cell line analogous to figure 2. AUROC curves are computed by iterating over silencer score quantile cutoffs from the set of all tested elements in 1 percentile increments.

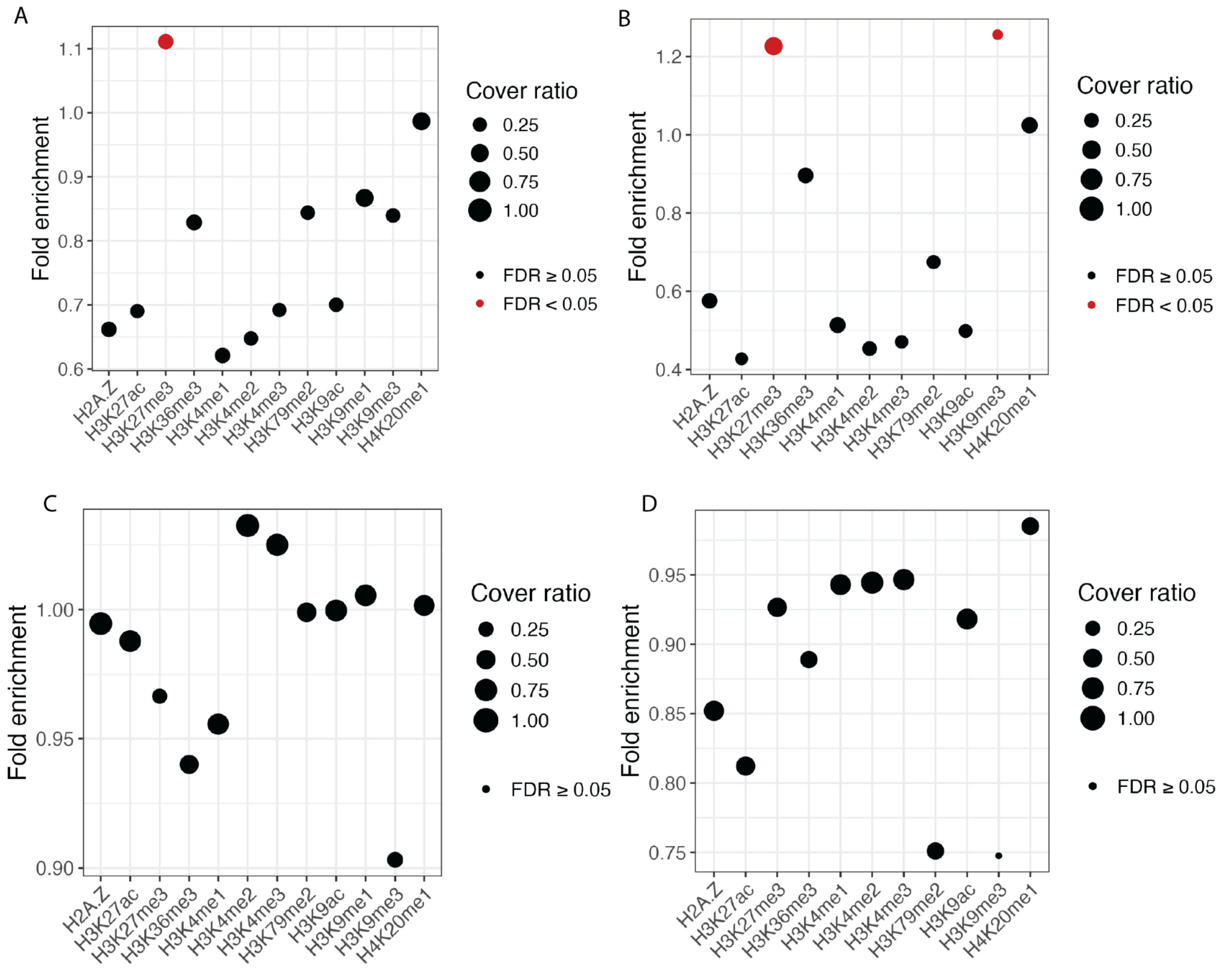


**Figure S2: Silencers are not enriched for any histone marks when considering FAIRE-seq peaks as background.** (**A**) Histone modifications enriched in ReSE K562 silencer elements using ENCODE FAIRE regions as background. Cover ratio denotes the fraction (0-1) of silencer elements overlapping the indicated peak set. Fold enrichment represents the cover ratio of the foreground silencers over the background cover ratio of non-silencer elements. P values are computed empirically by permutation test with 20,000 permutations of the background set and FDR corrected by Benjamini-Hochberg. Significant (FDR $q < 0.05$) enrichments are shown in red. (**B**) Histone modifications enriched in ReSE HepG2 silencer elements using ENCODE FAIRE regions as background, as in (**A**). (**C**) K562 silencers are not enriched for any histone mods when considering only elements that overlap K562 FAIRE-seq peaks. Cover ratio denotes the fraction (0-1) of silencer elements overlapping the indicated peak set. Fold enrichment represents the cover ratio of the foreground silencers over the background cover ratio of non-silencer elements. *P*-values were computed empirically by a permutation test with 20,000 permutations of the background set and FDR corrected by Benjamini-Hochberg. (**D**) Histone modification enrichments for ReSE HepG2 silencers, as in (**C**).

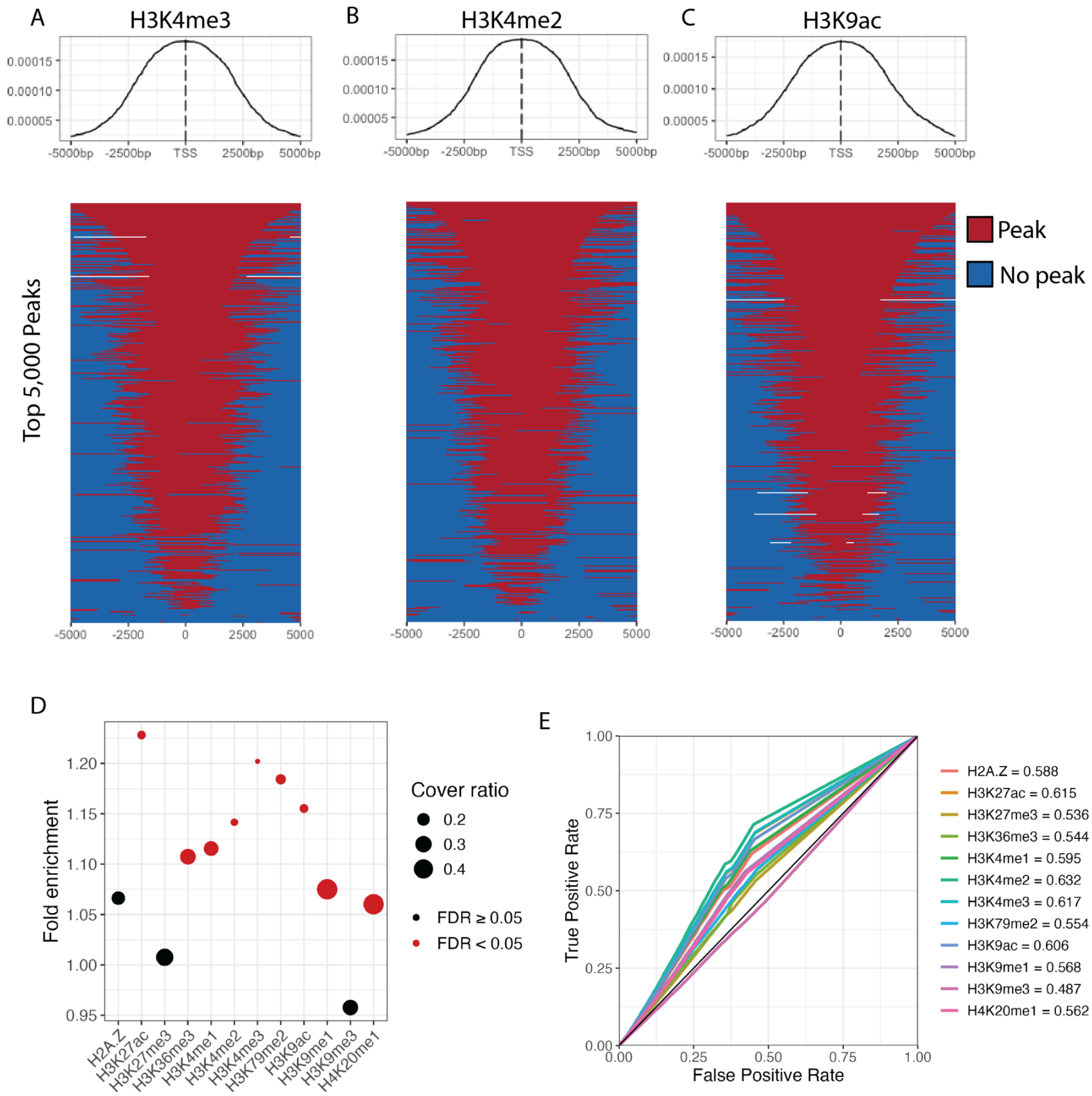


**Figure S3: Removing TSS-overlapping elements has a marginal effect on histone enrichments. (A-C)** Heatmaps showing the distributions of top 1000 peaks for (**a**) H3K4me3 , (**B**) H3K4me2 and (**C**) H3K9ac in K562 cells. (**D**) Enrichment values for K562 silencer elements after subtracting all tested elements within 5 kb of a TSS. Cover ratio denotes the fraction (0-1) of silencer elements overlapping the indicated peak set. Fold enrichment represents the cover ratio of the foreground silencers over the background cover ratio of non-silencer elements. *P*-values are computed empirically by a permutation test with 20,000 permutations of the background set and FDR corrected by Benjamini-Hochberg. Significant (FDR < 0.05) enrichments are shown in red. (**E**) AUROC curves for K562 silencers after subtracting all tested elements within 5 kb of a TSS.

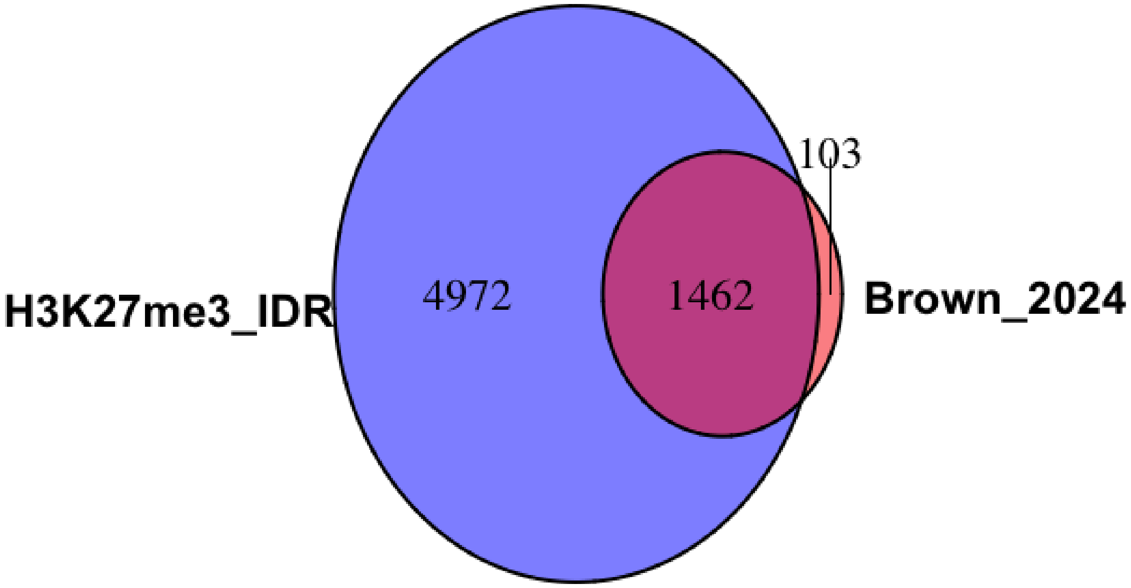


**Figure S4. Venn diagram of overlap between newly generated H3K27me3 peaks ("H3K27me3 IDR") with H3K27me3 peaks from Brown *et al.*, *Science Advances* (2024).** Elements were considered overlapping if they overlapped by at least 1 bp.

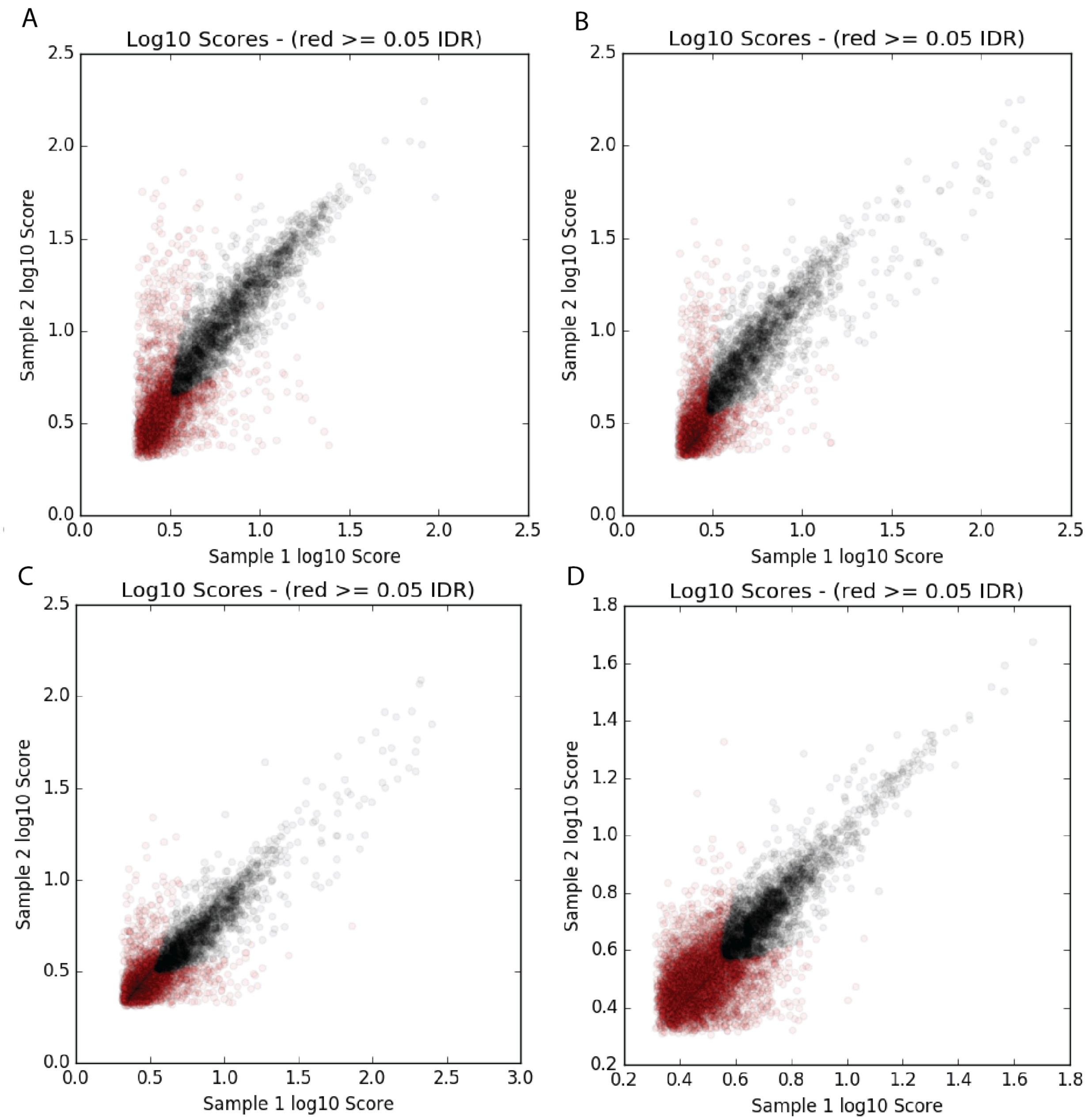


**Figure S5. Plots of IDR peak reproducibility between ChIP-seq replicates for each of the antibodies against histone modifications used in ChIP-Seq assays in *Drosophila* S2 cells in this study.** Scatterplots depict IDR for (**a**) Abcam 9051 H4K20me1, (**b**) Active Motif H4K20me1, (**c**) Thermo H4K20me1, and (**d**) Epicypher H3K27me3. Black dots represent reproducible peaks and red dots are not considered reproducible. All plots were generated by the IDR software.

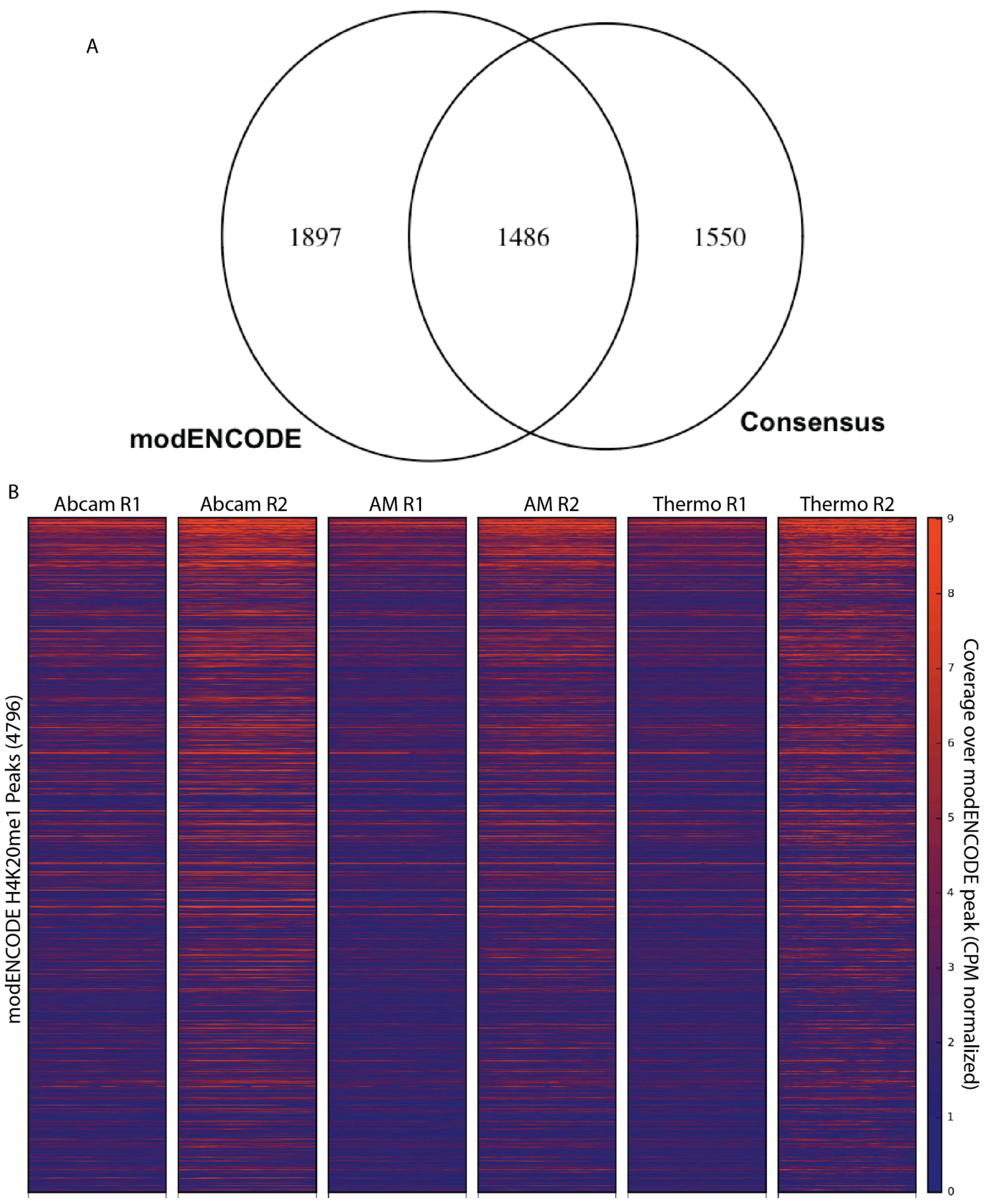
A
1897
1486
1550
modENCODE
Consensus
B
Abcam R1
Abcam R2
AM R1
AM R2
Thermo R1
Thermo R2
modENCODE H4K20me1 Peaks (4796)
Coverage over modENCODE peak (CPM normalized)
0
1
2
3
4
5
6
7
8
9

**Figure S6. S2 cell ChIP-seq corresponds to modENCODE ChIP-chip. A**) Venn diagram of overlaps (at least 1 bp) between modENCODE ChIP-chip peaks for H4K20me1 and the consensus set of H4K20me1 ChIP-Seq peaks generated in this study. **B**) Heatmap showing the CPM normalized coverage of H4K20me1 ChIP-seq reads over the 4796 modENCODE ChIP-chip peaks. Abcam 9051: Abcam ab9051; AM: Active Motif AB_2615074; Thermo: Thermo Fisher MA5-18067. Regions are sorted in descending order by statistical significance reported in the modENCODE ChIP-chip dataset from modENCODE Consortium, *Science* (2010).

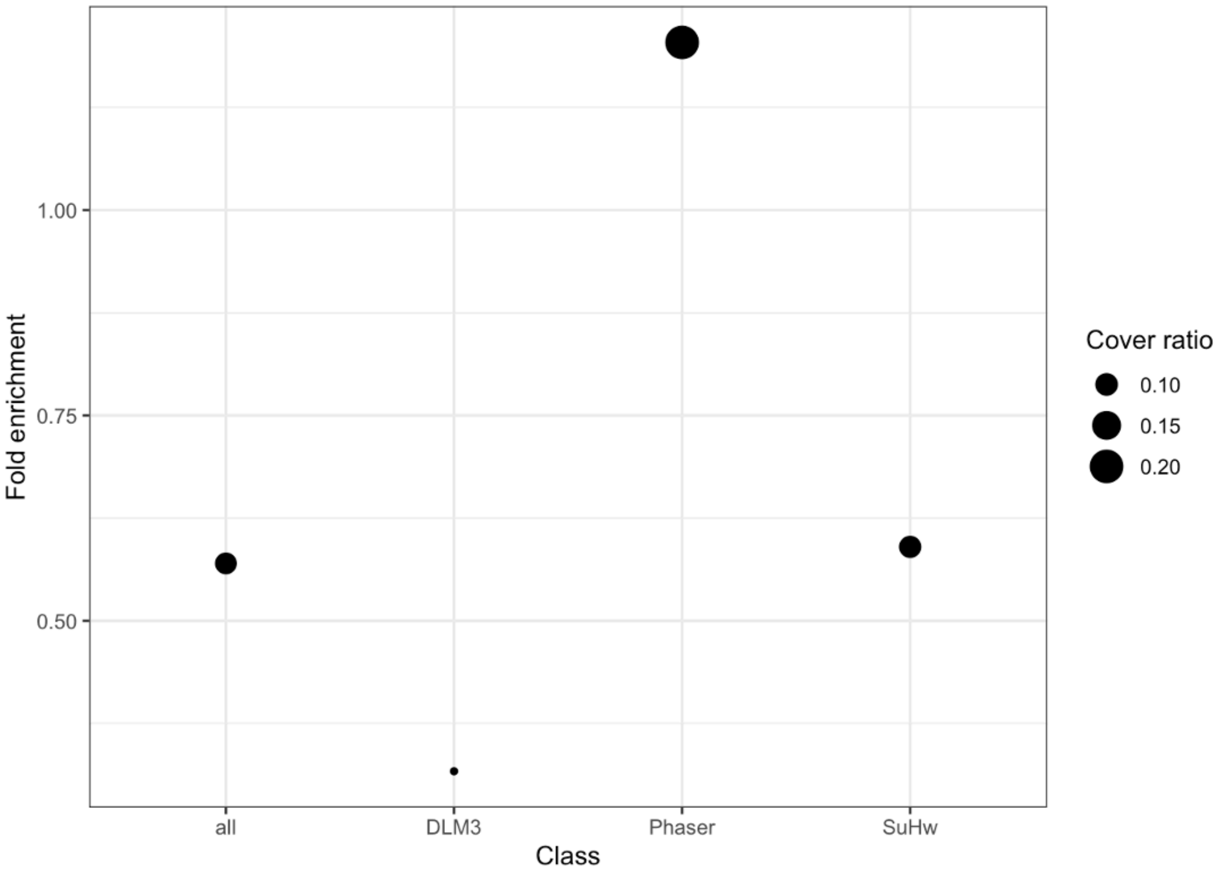


**Figure S7. H4K20me1 fold enrichment values for *Drosophila* S2 silencers as compared to GENRE background regions.** Background cover ratio is computed as the average of 100 GENRE backgrounds.

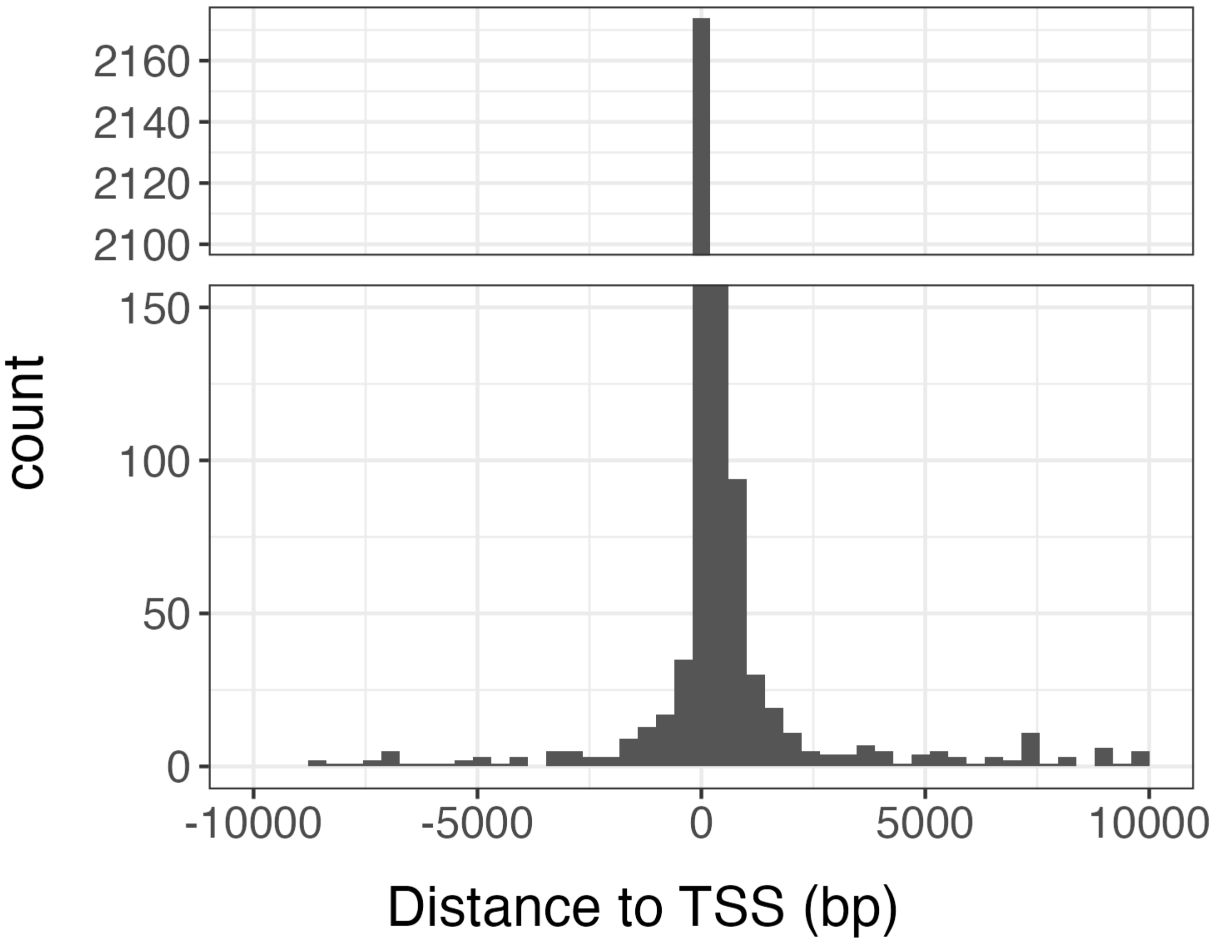


**Figure S8. H4K20me1 ChIP-Seq peaks in S2 cells are close to TSSs.** H4K20me1 consensus peaks were assigned to the closest TSS in Flybase annotation v.5.57.

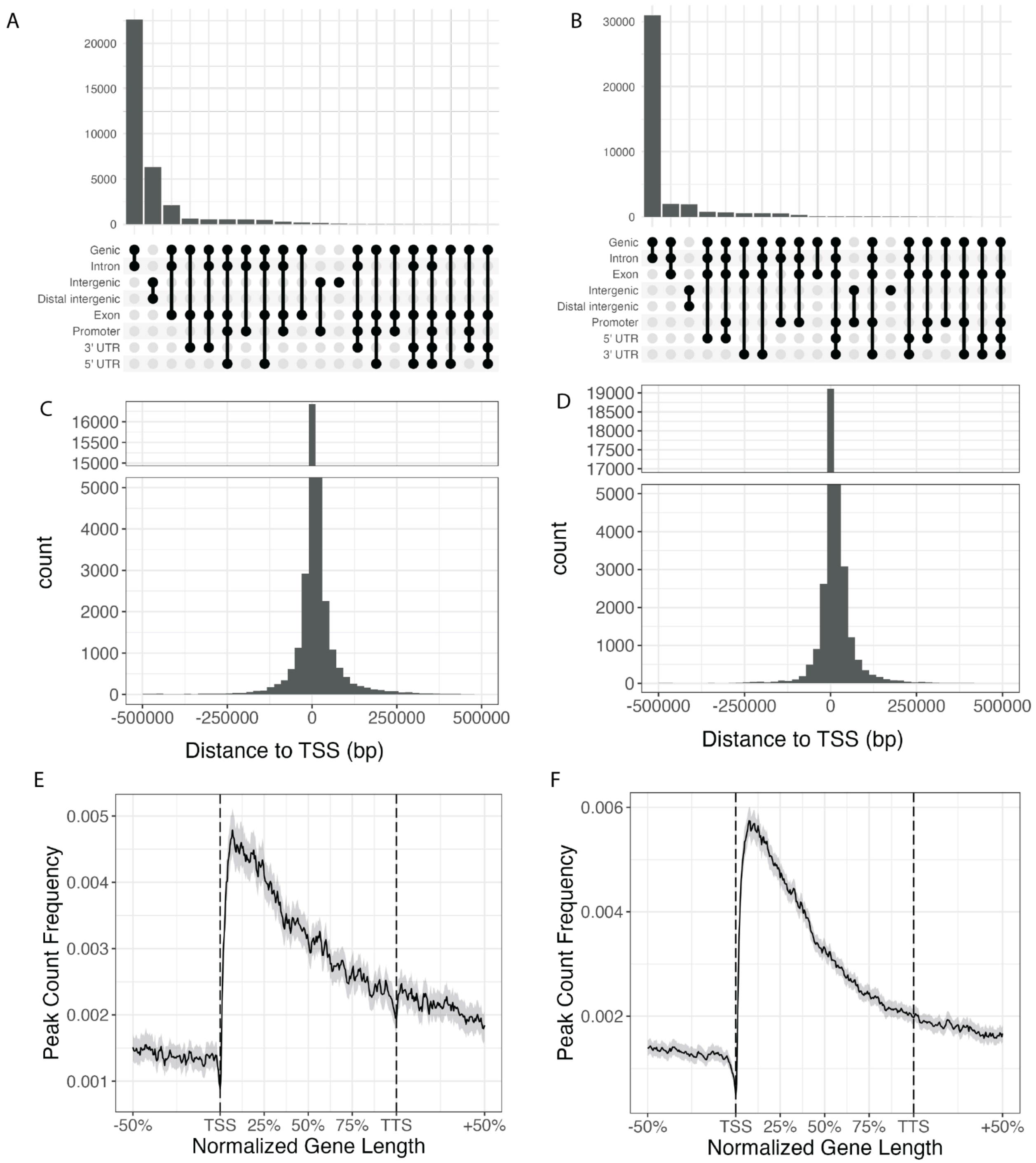


**Figure S9. H420me1 peaks in human cell lines are primarily genic.** (**A, B**) Upset plot of annotations for ENCODE H4K20me1 peaks from **A)** K562 and (**B**) HepG2 using Ensembl gene annotations for hg19. (**C, D**) Distribution of distances to TSS for ENCODE H4K20me1 peaks in (**C**) K562 and (**D**) HepG2. (**E, F**) Aggregate peak profile of ENCODE H4K20me1 peaks in (**E**) K562 and (**F**) HepG2 over genes.

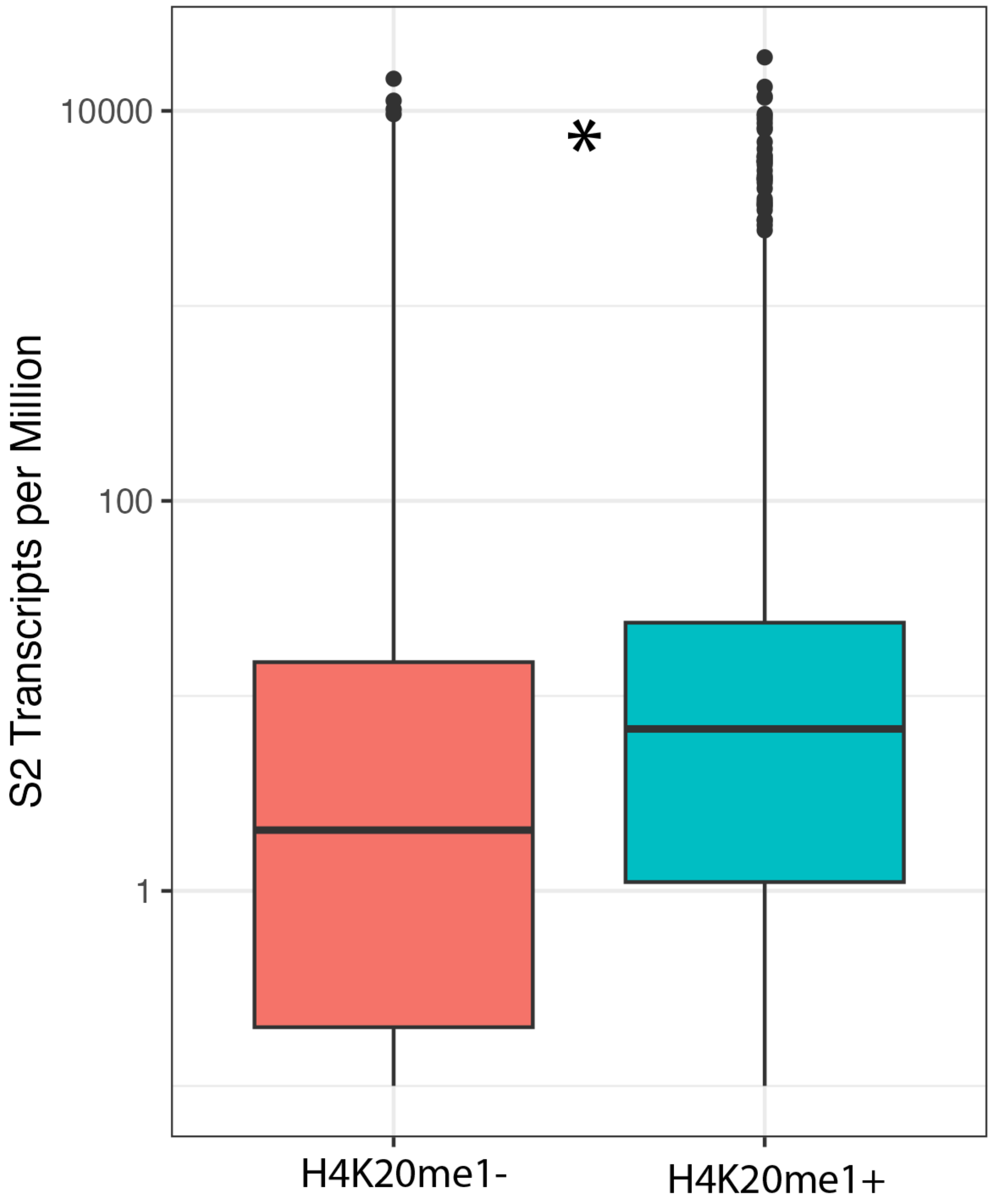


**Figure S10. Genes marked by H4K20me1 are more highly expressed in S2 cells.** RNA-seq TPM values are from Klonaros et al. (2023) G3. * : $P < 0.05$ by Wilcoxon rank-sum test.

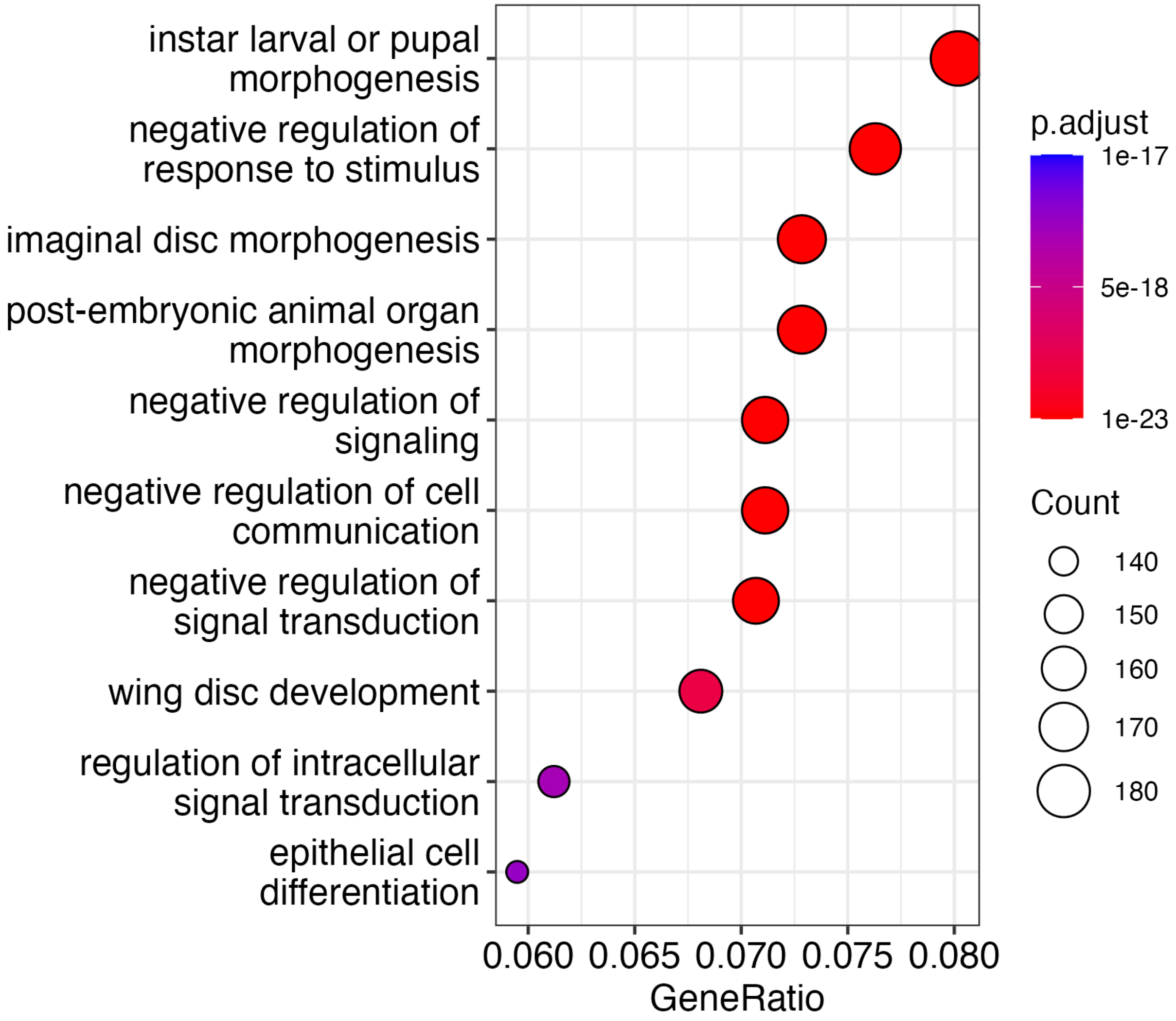


**Figure S11. H4K20me1 marks developmentally active genes.** Top 10 enriched GO annotations among genes that are covered by H4K20me1 peaks in S2 cells. GeneRatio refers to the proportion of genes in the list that have the indicated GO annotation.

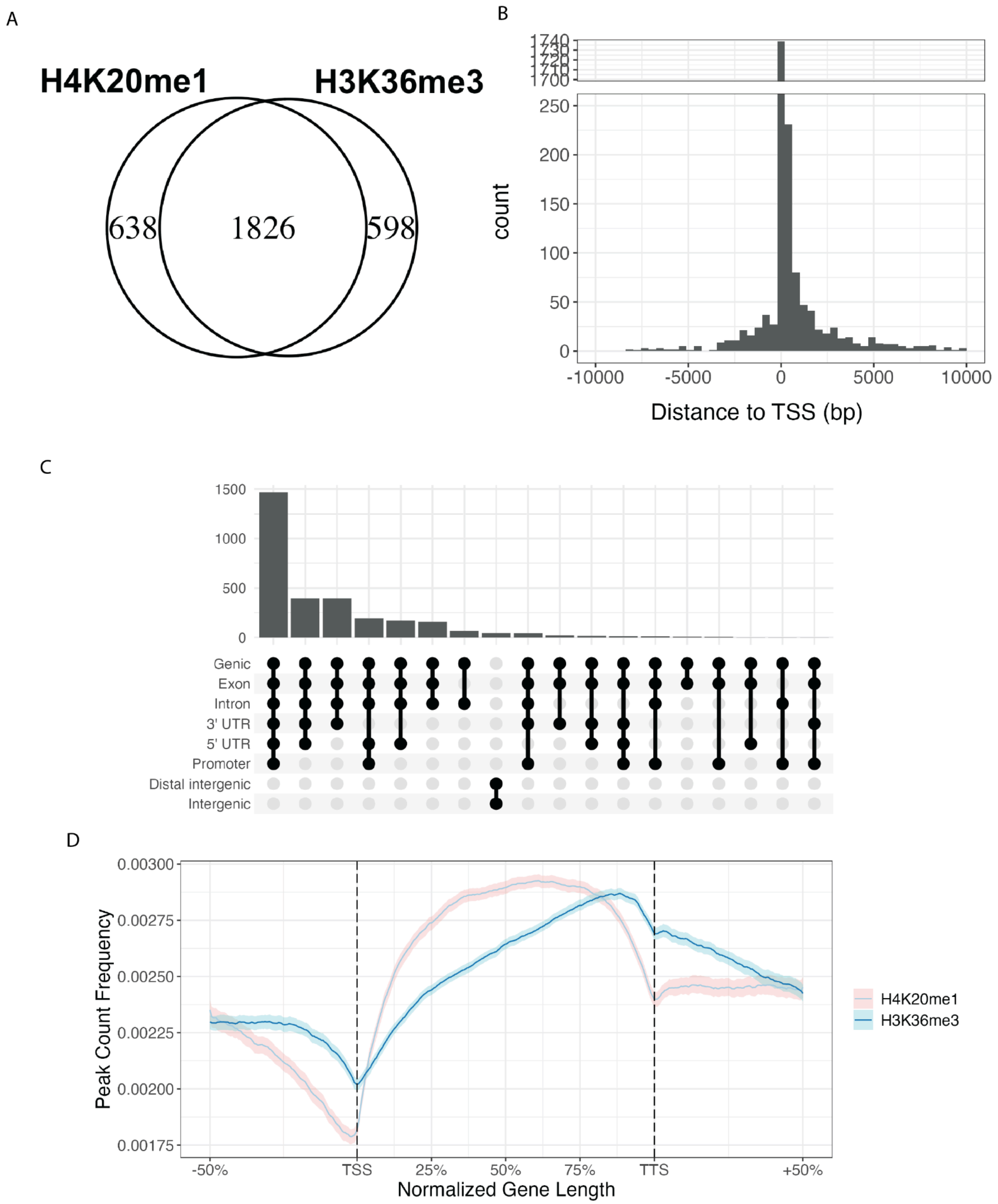


**Figure S12. H3K36me3 has a distribution similar to that of H4K20me1 in *Drosophila* S2 cells.** (**A**) Venn diagram of peak overlap between S2 H4K20me1 peaks and H3K36me3 peaks. (**B**) Distribution of distances to nearest TSS for H3K36me3 peaks. (**C**) Upset plot of genomic annotations for H3K36me3 peaks. (**D**) Aggregate peak profile of H3K36me3 over genes. H3K36me3 peaks are from Brown *et al.*, *Science Advances* (2024).

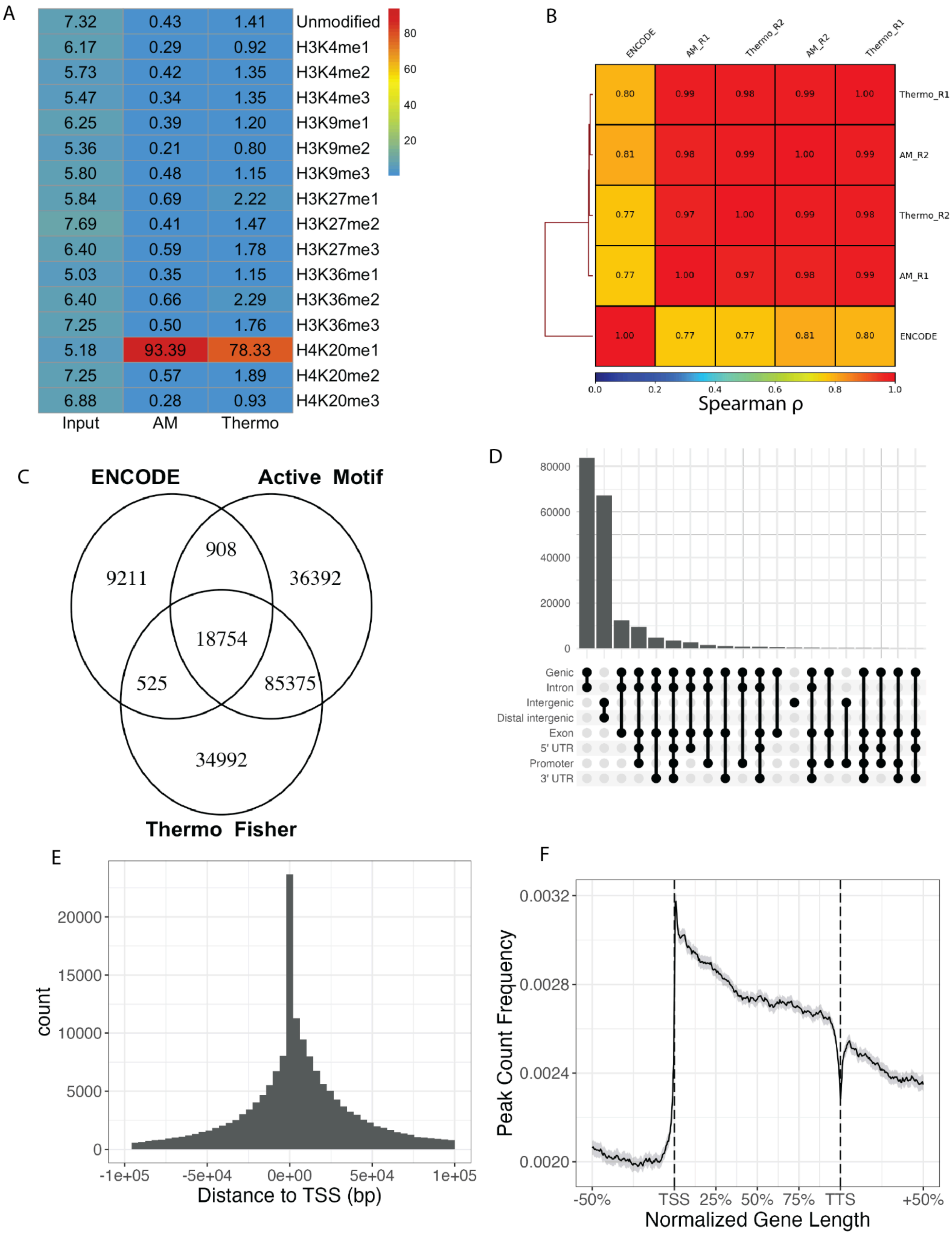

A
B
C
ENCODE
Active Motif
Thermo Fisher
D
E
F
Spearman ρ
Input
AM
Thermo
Distance to TSS (bp)
count
Normalized Gene Length
Peak Count Frequency

**Figure S13: K562 ChIP-seq libraries show consistency with ENCODE data.** (A) Summary of antibody specificity as measured by SNAP-ChIP spike-in mononucleosomes. Columns represent individual ChIP-seq libraries, rows represent all histone modifications present in the panel. Values denote percentages of barcode reads mapped to each modification within a library. (B) Matrix of Spearman correlation between H4K20me1 ChIP-seq libraries across all genomic bins. (C) Venn diagram showing the number of overlapping peaks across H4K20me1 datasets. ENCODE peak set was generated by ENCODE Consortium using Abcam ab9051. (D) Upset plot showing the genomic annotations of consensus H4K20me1 peaks from UCSC annotation. (E) Histogram of distances to nearest TSS for H4K20me1 consensus peaks. (F) Aggregate profile of H4K20me1 consensus peak coverage over genes.

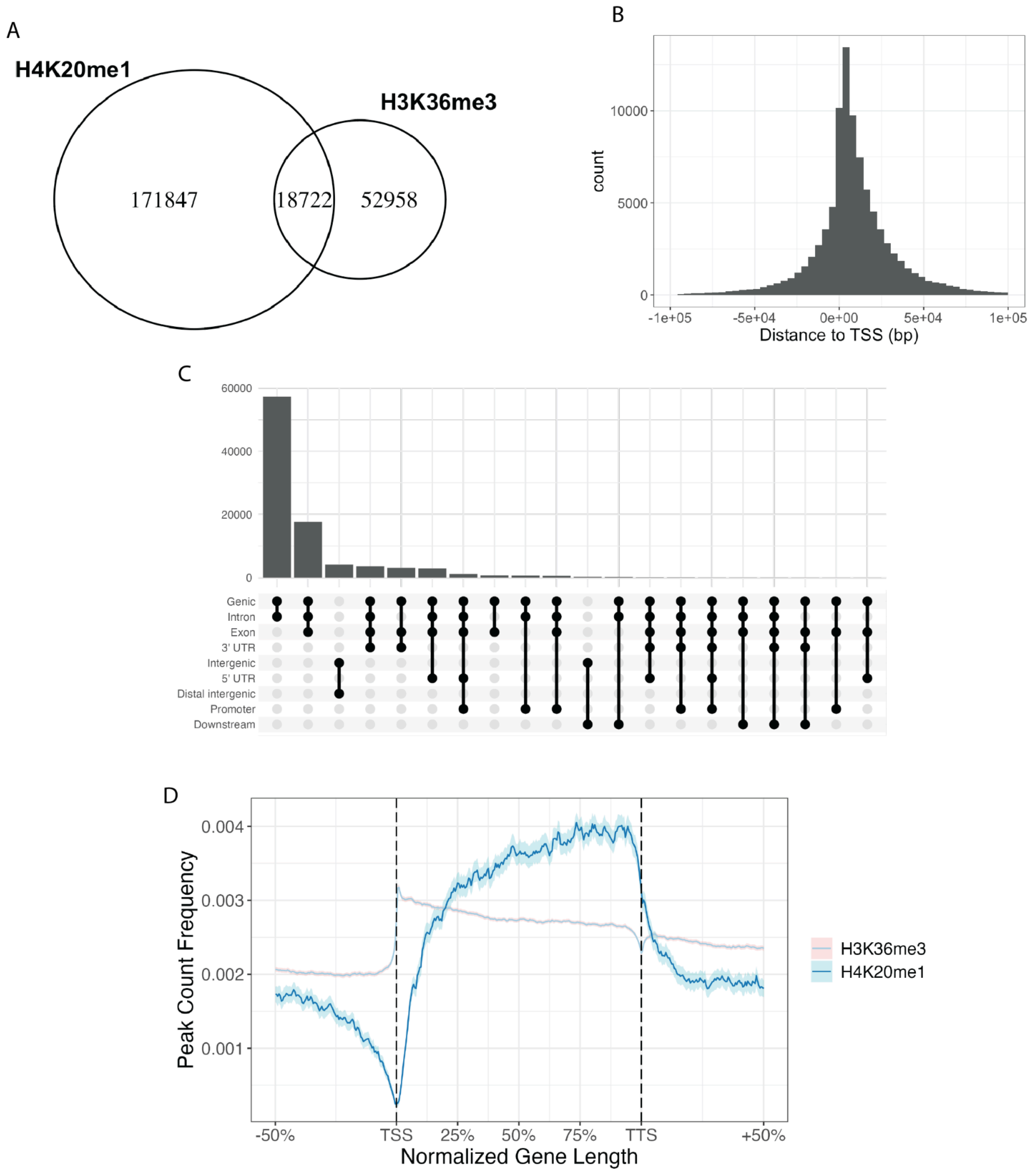


**Figure S14. H3K36me3 has a distribution similar to that of H4K20me1 in K562 cells.** (**A**) Venn diagram of peak overlap between K562 H4K20me1 peaks and ENCODE H3K36me3 peaks. (**B**) Distribution of distances to nearest TSS for ENCODE H3K36me3 peaks. (**C**) Upset plot of genomic annotations for ENCODE H3K36me3 peaks. (**D**) Aggregate peak profile of ENCODE H3K36me3 over genes.

| Sample ID | # Reads (paired) | Mean Quality Score | % Bases ≥ 30 | % mapped | % duplicate | # reads mapq > 30 |
|---|---|---|---|---|---|---|
| Input-R1 | 26575237 | 39.37 | 96.91 | 97.48 | 0.115781 | 33408296 |
| Input-R2 | 30796771 | 39.31 | 96.65 | 97.93 | 0.1252 | 38518382 |
| H4K20me1-Abcam9051-R1 | 28267240 | 39.31 | 96.62 | 96.96 | 0.099941 | 37835128 |
| H4K20me1-Abcam9051-R2 | 24183931 | 38.79 | 94.17 | 95.4 | 0.098904 | 29475716 |
| H4K20me1-AM-R1 | 22135191 | 39.08 | 95.59 | 85.96 | 0.137128 | 25548658 |
| H4K20me1-AM-R2 | 14984036 | 39.06 | 95.48 | 86.00 | 0.135871 | 17451860 |
| H4K20me1-Thermo-R1 | 30608549 | 39.06 | 95.46 | 86.02 | 0.139605 | 35484084 |
| H4K20me1-Thermo-R2 | 5788272 | 38.89 | 94.66 | 82.00 | 0.136195 | 6323372 |
| H3K27me3-R1 | 30272558 | 39.4 | 97.06 | 97.00 | 0.094009 | 42923906 |
| H3K27me3-R2 | 25704490 | 39.09 | 95.59 | 96.88 | 0.089931 | 35214128 |

**Table S1. Sequencing statistics for ChIP-seq and corresponding input libraries.** All reads are paired-end 2 x 150 bp reads sequenced in the same run. Duplication percentage was calculated by Picard MarkDuplicates. Percentage mapped and mapq scores are given by Bowtie2 using dm3 as the reference genome.

| Sample ID | MACS2 peaks | FRiP | MACS2 covered bases | IDR of MACS2 peaks | IDR bases covered | Percent of peaks passing IDR |
|---|---|---|---|---|---|---|
| H4K20me1-Abcam9051-R1 | 6162 | 0.3978 | 19100895 | | | |
| H4K20me1-Abcam9051-R2 | 5774 | 0.5734 | 23281427 | 4958 | 28245689 | 48.5 |
| H4K20me1-AM-R1 | 4538 | 0.2779 | 14008740 | | | |
| H4K20me1-AM-R2 | 4696 | 0.4239 | 17935957 | 4104 | 24127849 | 50.1 |
| H4K20me1-Thermo-R1 | 5032 | 0.3130 | 16712011 | | | |
| H4K20me1-Thermo-R2 | 4695 | 0.4088 | 16712011 | 4301 | 25360642 | 44.1 |
| H3K27me3-R1 | 10162 | 0.6355 | 42640439 | | | |
| H3K27me3-R2 | 10649 | 0.7045 | 42675048 | 7142 | 54446766 | 36.7 |

**Table S2. Peak calling statistics from *Drosophila* H4K20me1 ChIP-seq libraries.** MACS2 peaks for each library were called using the default q value cutoff of 0.05, whereas idr peaks were called using a more relaxed *P*-value cutoff of 0.01 for each replicate. Fraction of reads in peaks (FRiP) was computed using the $q < 0.05$ peaks for each library individually. IDR peaks are only listed for replicate 1 but apply to replicates 1 and 2 together. Abcam 9051: Abcam ab9051; AM: Active Motif AB_2615074; Thermo: Thermo Fisher MA5-18067.

| Sample ID | # Reads (paired) | % Bases ≥ 30 | % mapped | # mapped reads | % duplicate |
|---|---|---|---|---|---|
| Input-R1 | 83,099,036 | 100% | 96.94 | 157246482 | 0.062601 |
| Input-R2 | 96,699,107 | 100% | 96.90 | 183138218 | 0.071756 |
| H4K20me1-Active-Motif-R1 | 155,867,997 | 100% | 78.38 | 230920478 | 0.153481 |
| H4K20me1-Active-Motif-R2 | 163,675,747 | 100% | 75.53 | 230326278 | 0.163276 |
| H4K20me1-Thermo-R1 | 172,192,983 | 100% | 68.48 | 218986122 | 0.114591 |
| H4K20me1-Thermo-R2 | 225,696,067 | 100% | 70.34 | 298224500 | 0.140748 |
| H3K27me3-R1 | 178,181,080 | 100% | 84.22 | 288733690 | 0.115741 |
| H3K27me3-R2 | 104,225,510 | 100% | 77.37 | 154298964 | 0.113486 |

**Table S3. Sequencing statistics for K562 ChIP-seq and corresponding input libraries.** All reads are paired-end 2 x 150 bp reads sequenced in the same run. Duplication percentage was calculated by Picard MarkDuplicates. Percentage mapped and mapq scores are given by Bowtie2 using hg19 as the reference genome.

| Sample ID | MACS2 peaks | FRiP | MACS2 covered bases | IDR of MACS2 peaks | IDR bases covered | Percent of peaks passing IDR |
|---|---|---|---|---|---|---|
| H4K20me1-Active-Motif-R1 | 258544 | 0.365041116 | 228674421 | | | |
| H4K20me1-Active-Motif-R2 | 180801 | 0.3172867666 | 225381756 | 264921 | 529965292 | 9.5 |
| H4K20me1-Thermo-R1 | 215859 | 0.3513906009 | 227108228 | | | |
| H4K20me1-Thermo-R2 | 217804 | 0.394151225 | 249645072 | 268677 | 498290978 | 13.2 |
| H3K27me3-R1 | 254039 | 0.4531146227 | 313662034 | | | |
| H3K27me3-R2 | 287150 | 0.4750954582 | 318716561 | 291796 | 570922858 | 28.4 |

**Table S4. Peak calling statistics from K562 H4K20me1 ChIP-seq libraries.** MACS2 peaks for each library were called using the default q value cutoff of 0.05, whereas idr peaks were called using a more relaxed *P*-value cutoff of 0.01 for each replicate. Fraction of reads in peaks (FRiP) was computed using the q < 0.05 peaks for each library individually. IDR peaks are only listed for replicate 1 but apply to replicates 1 and 2 together. AM: Active Motif AB_2615074; Thermo: Thermo Fisher MA5-18067.
