## Supplementary material for "Histone H4 lysine 20 monomethylation is not a mark of transcriptional silencers": Table S5, Supplemental methods, Supplementary references

| Modification | K562 UCSC accession | HepG2 UCSC accession |
|---|---|---|
| H3K4me3 | wgEncodeEH000048 | wgEncodeEH000095 |
| H3K36me3 | wgEncodeEH000045 | wgEncodeEH000081 |
| H3K4me1 | wgEncodeEH000046 | wgEncodeEH001749 |
| H3K27me3 | wgEncodeEH000044 | wgEncodeEH001023 |
| H3K9ac | wgEncodeEH000049 | wgEncodeEH000083 |
| H2A.Z | wgEncodeEH001038 | wgEncodeEH001022 |
| H4K20me1 | wgEncodeEH000051 | wgEncodeEH000096 |
| H3K9me3 | wgEncodeEH001040 | wgEncodeEH003087 |
| H3K4me2 | wgEncodeEH000047 | wgEncodeEH000082 |
| H3K27ac | wgEncodeEH000043 | wgEncodeEH000094 |
| H3K79me2 | wgEncodeEH001039 | wgEncodeEH001024 |
| H3K9me1 | wgEncodeEH000050 | N/A |

**Table S5: UCSC Genome Browser accession numbers for K562 and HepG2 histone ChIP-seq peaks.**

### Supplemental methods

#### S2 cell collection and chromatin shearing

Cells were harvested by use of a plastic cell scraper, tapping the flask, and repeated pipetting of media. Approximately 2 x $10^7$ cells were used per chromatin immunoprecipitation (ChIP) reaction, although the final amount was determined by micrograms of DNA and not input cell number as this reduced variability from automated cell counting and variable efficiency of sonication. Cells were pelleted by centrifugation at 1000 x g for 5 minutes and washed twice with 1X PBS (Gibco 10010023). They were then crosslinked with 1% formaldehyde (Electron Microscopy Science 15710) in PBS at room temperature for 10 minutes with shaking. Crosslinking was quenched by the addition of 2.5 M glycine to a final concentration of 125 mM and incubated at room temperature for 5 minutes. Cells were washed twice with cold PBS and finally resuspended in PBS + complete protease inhibitor cocktail EDTA-free (Millipore Sigma 11873580001) and divided into aliquots of ~1 x $10^8$ cells in microcentrifuge tubes. These were pelleted again at 1000 x g for 5 minutes, the supernatant was aspirated off, and the pellets were frozen at -80 °C for future processing. Two replicates were collected and fixed in different batches on different days.

To shear chromatin, cell pellets were thawed on ice and resuspended in cell lysis buffer and incubated for 10 minutes. Samples were centrifuged at 4000 x g for 5 minutes and then resuspended in 1mL nuclear lysis buffer. Samples were sonicated in a Bioruptor Plus (Diagenode) at high power for 30 seconds on and 30 seconds off. The number of cycles was chosen to fragment DNA with a peak of fragments ~200 - 600 bp. In our case, 60 cycles were needed, but the exact number will vary depending on many factors and should be determined empirically by testing many iterations. To measure DNA concentration, a 100 µL aliquot was decrosslinked by adding 38 uL decrosslinking buffer (2 M NaCl, 0.1M EDTA, 0.4M Tris pH 7.5 [1]) and incubating overnight at 65 °C followed by incubation with 2 µL proteinase K (Thermo

Fisher 25530049) at 50 °C for 2 hours. DNA was extracted by phenol-chloroform purification and then precipitated with ethanol and linear acrylamide (Thermo Fisher AM9520) as a carrier. DNA pellets were reconstituted in 100 µL water and quantified by Qubit HS dsDNA kits (Invitrogen Q33231).

**ChIP-seq**

To evaluate antibody specificity, 10 µL SNAP-ChIP K-MetStat mononucleosomes (Epicypher 19-1001) were spiked into each replicate chromatin prep before aliquoting into individual ChIP reactions. 100 µL input chromatin (“Input”) was reserved and sequenced as background. To dilute the concentration of sodium dodecyl sulfate (SDS) in the nuclear lysis buffer prior to ChIP, sonicated chromatin was diluted 1:5 with IP dilution buffer (16.7 mM Tris pH 8.0, 1.2 mM EDTA, 167 mM NaCl, 1.1% Triton X-100, 0.01% SDS) [1]. 100 µg chromatin as measured from the decrosslinked sample was aliquoted to each ChIP reaction.

ChIP was performed using several different antibodies against H4K20me1 to check for consistency of ChIP signal and ensure high quality data. The H4K20me1 antibodies used were: Abcam ab9051, used widely by ENCODE and modENCODE as well as [2] Active Motif AB_2615074, which has not been used in a published ChIP-seq study to our knowledge; and Thermo Fisher MA5-18067, which was recently used in CUT&RUN experiments in *Drosophila* [3]. As a positive control, ChIP was performed in parallel for H3K27me3 using a well characterized H3K27me3 antibody (Epicypher 13-0055).

Chromatin was first pre-cleared by incubation with 20 uL unblocked Magna ChIP Protein A+G beads (Sigma Aldrich 16-663). Beads were washed once in IP dilution buffer for 2 hours with rotation before adding to chromatin. The beads were then immobilized with a magnet and the precleared chromatin was moved to new tubes and then incubated with 5ug antibody (see above) overnight at 4C with rotation. In parallel, beads for ChIP were blocked overnight in 1 mg/mL BSA (NEB B9200S) in IP dilution buffer at 4C. Beads were washed once with IP dilution buffer before 20uL beads were added to each sample and then incubated for 2 hours at 4C. Washes and elution were performed as previously described [1]. Briefly, beads were washed 6 times with low salt buffer (0.1% SDS, 1% Triton X-100, 2 mM EDTA, 20 mM Tris pH 8.0, 150 mM NaCl), once with high salt buffer (0.1% SDS, 1% Triton X-100, 2 mM EDTA, 20 mM Tris pH 8.0, 500 mM NaCl), and once with TE pH 8.0 (Invitrogen AM9849) before elution with 250 µL elution buffer (0.1M NaHC03, 1% SDS). Eluted DNA was decrosslinked by overnight incubation with decrosslinking buffer at 65 °C and purified using Monarch PCR and DNA Cleanup Kit (New England Biolab T1030S). DNA was quantified using Qubit HS dsDNA and High Sensitivity D5000 DNA ScreenTape assays (Agilent).

**Generation and QC of sequencing libraries**

Sequencing libraries were generated using NEBNext Ultra II (E7645S) with NEBNext multiplex oligos for Illumina (E7335S). The number of PCR cycles was selected according to manufacturer's recommendations for the mass of DNA in each library. Input, H4K20me1 libraries using Abcam 9051 antibodies, and H3K27me3 libraries were amplified with 3 cycles of PCR, while H4K20me1 libraries using the Active Motif or Thermo antibodies required 6 cycles of PCR. QC of sequencing libraries was performed using Qubit High Sensitivity dsDNA assays for DNA concentration and High Sensitivity D1000 and D5000 DNA ScreenTape assays (Agilent) for fragment length distributions. Since the libraries generated using the Active Motif or Thermo H4K20me1 antibodies skewed towards larger fragment sizes, we size-selected them with SPRISelect beads (Beckman Coulter B23317) prior to sequencing, using a right-side size selection with 0.4x bead ratio to remove large fragments. Libraries were paired-end sequenced 2 x 150bp on an Illumina HiSeq 2500. Sequencing statistics are provided in Additional file 4: Table S3.

**ChIP-seq data processing**

Trim galore (v. 0.6.6) [4] was used to remove adaptor sequences and bases with sequencing quality below 30. Reads were then mapped to dm3 using bowtie2 (v. 2.5.1) [5] with the settings "--sensitive-local --no-discordant --no-mixed". PCR duplicates were called using Picard MarkDuplicates. Libraries were filtered for alignments above mapq 30. Peak calling was performed using MACS2 [6] (v. 2.1.1.20160309) with the corresponding input reads as control and the settings "--broad -g dm -f BAMPE". Following ENCODE guidelines [7], we used the irreproducible discovery rate (IDR) framework [8] to identify peaks that are consistent between the two replicate samples that used the same antibody. For calling peaks found to be reproducible by IDR analysis, a more relaxed threshold of "-p 1e3" was used to call peaks in each library before running IDR [8] (v. 2.0.2). A summary of peak counts is provided in Additional file 5: Table S4. High confidence "consensus" H4K20me1 peaks used for downstream analyses were determined by concatenating the sets of IDR reproducible peaks and retaining only those peaks that were common across all three peak sets from the three H4K20me1 antibodies that we used. Consensus peaks were then merged to remove redundancies and  overlapping peaks were collapsed into continuous regions.

Peaks were annotated using the annotatePeak function of ChIPSeeker (v.1.38.0) [9] with the reference annotation packages TxDb.Hsapiens.UCSC.hg19.knownGene (v.3.2.2) for humans and Flybase (v5.7) for *Drosophila*. Bed files of H4K20me1 ChIP-seq peaks in K562 and HepG2 used for genomic annotation were downloaded from ENCODE (accessions ENCFF139CKE and ENCFF869HGO respectively). The TSS region was set to 250bp downstream and 50bp upstream for *Drosophila* and 500bp downstream and 500bp upstream for human. Names of H4K20me1 intersecting genes were determined according to org.Dm.eg.db (v.3.19.0). Gene Ontology (GO) term analysis was performed using the clusterProfiler (v.4.10.1) [10] package.
